# Naturally ornate RNA-only complexes revealed by cryo-EM

**DOI:** 10.1101/2024.12.08.627333

**Authors:** Rachael C. Kretsch, Yuan Wu, Svetlana A. Shabalina, Hyunbin Lee, Grace Nye, Eugene V. Koonin, Alex Gao, Wah Chiu, Rhiju Das

## Abstract

Myriad families of natural RNAs have been proposed, but not yet experimentally shown, to form biologically important structures. Here we report three-dimensional structures of three large ornate bacterial RNAs using cryogenic electron microscopy at resolutions of 2.9-3.1 Å. Without precedent among previously characterized natural RNA molecules, Giant, Ornate, Lake- and Lactobacillales-Derived (GOLLD), Rumen-Originating, Ornate, Large (ROOL), and Ornate Large Extremophilic (OLE) RNAs form homo-oligomeric complexes whose stoichiometries are retained at concentrations lower than expected in the cell. OLE RNA forms a dimeric complex with long co-axial pipes spanning two monomers. Both GOLLD and ROOL form distinct RNA-only multimeric nanocages with diameters larger than the ribosome. Extensive intra- and intermolecular A-minor interactions, kissing loops, an unusual A-A helix, and other interactions stabilize the three complexes. Sequence covariation analysis of these large RNAs reveals evolutionary conservation of intermolecular interactions, supporting the biological importance of large, ornate RNA quaternary structures that can assemble without any involvement of proteins.

---

The importance of non-coding RNAs across biology is increasingly appreciated, yet only a handful have been functionally characterized, with studies revealing sophisticated catalytic and sensory functions in some cases^1–5^. Bacteria, archaea and their viruses are hypothesized to possess numerous diverse and complex ncRNAs, but most of these have not been thoroughly studied^6–9^. Furthermore, there is a conspicuous shortage of data on the three-dimensional structures of RNA molecules. Out of more than 4,000 RNA classes in the RNA Families (RFAM) database 15.0, only 143 have experimentally resolved tertiary structures^10^. For many of the remaining cases, it appears likely that structural characterization will depend on reconstitution of the RNA with small molecule, protein, or nucleic acid partners, which are unknown in most cases.

The Breaker laboratory and collaborators have previously described three classes of bacterial and phage RNAs for which covariance analysis of genomic and metagenomic sequences revealed secondary structures so extensive and elaborate that epithets ‘ornate’, and ‘giant’ or ‘large’ were included in their names: Giant, Ornate, Lake- and Lactobacillales-Derived (GOLLD) RNA^8^; Rumen-Originating, Ornate, Large (ROOL) RNA^6,11^ (concomitantly reported in ref.^12^); and Ornate Large Extremophilic (OLE) RNA^7^. The functions of all these three classes of large RNAs remain poorly understood.

Here, using cryogenic electron microscopy (cryo-EM), we show that OLE, ROOL, and GOLLD all form atomically ordered three-dimensional structures. Unexpectedly, all three structures are stabilized not by proteins but by other copies of the same RNA molecule in ornate quaternary assemblies with numerous intermolecular bridges, a phenomenon not previously observed for natural RNA molecules^13^.

## OLE forms an RNA-only dimer presenting protein binding sites

OLE is a class of large RNAs with an ornate secondary structure conserved throughout evolution^7^. OLE is mainly found in extremophilic bacteria, and experimental characterization in *Halalkalibacterium halodurans* demonstrated OLE’s involvement in integrating energy availability, metal ion homeostasis, and drug treatment to mediate cellular adaptation, although the underlying molecular mechanisms remain unknown^7,14–16^. Cellular localization to the membrane, binding to at least six protein partners^15,17–22^, and evidence of alternative secondary structures^17^ suggested that OLE is unlikely to form a well-defined RNA-only three-dimensional structure.

However, our study showed that the 577-nucleotide (nt) OLE RNA from *Clostridium acetobutylicum*^7,23^ formed distinct, compact particles that were clearly visible in cryo-EM images (**Fig. 1A**). A 2.9 Å resolution 3D map could be reconstructed with two-fold imposed symmetry (**Extended Data** Figure 1). A model of the each chain has been built for 308 nt in the OLE 5′ region, with Q-scores^24^ exceeding that expected at this resolution (**Fig. 1B, Extended Data** Fig. 1E, and **Supplemental Movie 1**). This is in contrast with an RNA family with similar size to the OLE 5′ region, the HNH Endonuclease-Associated RNA and ORF (HEARO)^8^, which is known to form a defined RNA structure that is involved in DNA nickase activity when bound to the protein IsrB^25^; our cryo-EM data indicate that the 343-nt HEARO RNA from *Limnospira maxima* is disordered in the absence of the protein (**Extended Data** Figure 2), whereas OLE RNA is ordered without protein.

**Figure 1.**
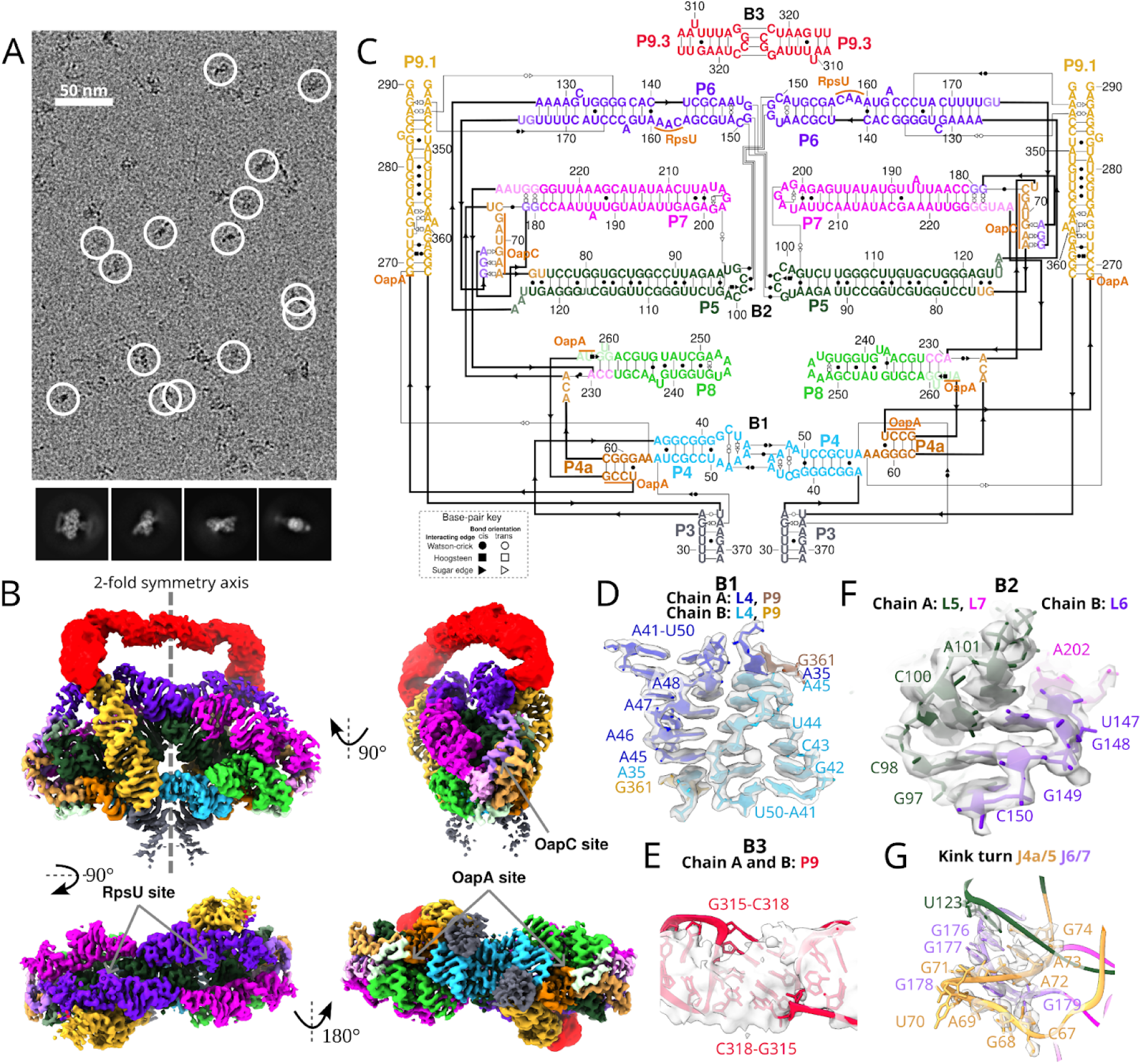
Structure of OLE homo-dimer. (**A**) Representative micrograph, particles selected for reconstruction are circled in white, and below are 2D class averages. (**B**) The cryo-EM reconstruction of the OLE dimer. The top, left image depicts the separation of the two chains, one chain on the right and the other on the left. Each domain of both chains is colored. To aid visualization, the flexible P9.3 domain (red) is displayed at a different contour and sharpening. The proposed binding sites of previously described proteins (RpsU, OapC, OapA) are labeled. (**C**) Secondary structure of OLE dimer labeling the domains colored like the cryo-EM map in (B). (**D-F**) The three intermolecular bridge interactions (B1, B2 and B3) colored by domain. In (D) the domain coloring is darkened for chain A to differentiate the chains. (**G**) Kink-turn motif that may bind the OapC protein, identical for each monomer.

Our OLE dimer map shows that it is organized as a series of parallel A-form helices, like a bundle of pipes. The exterior ends of these pipes from each chain are interconnected into a five-way junction, with a secondary structure agreeing with the previously proposed one for the observed domain with stems P3 to P9.3^7,15^ (**Fig. 1C**; hereafter, paired stems, hairpin loops, and joining linkers are labeled ‘P’, ‘L’, and ‘J’, respectively, following conventional RNA nomenclature). An unusual but highly conserved symmetric interaction comprised of four A-A base pairs between two chains (L4, **Fig. 1D**), intermolecular base-pairing and stacking interactions connecting L5, L6, and L7 (**Fig. 1E**), and a kissing loop (L9.3, **Fig. 1F**) ‘weld’ the pipes together in the middle of the complex. Hereafter we denote these intermolecular interactions ‘bridges’ B1-B3, as used in ribosome nomenclature^26^. Beyond the resolved 5′ region, other conserved parts of OLE were not resolved in the structure, suggesting flexibility.

Surprisingly, regions of OLE previously thought to adopt alternative structures upon protein binding are clearly resolved and solvent accessible, suggesting that proteins may bind the OLE dimer in the pre-formed RNA conformation observed here. Our cryo-EM data show that these proteins are not required for the folding of the 5′ domain of OLE and the RNA structure itself may play a crucial role in organizing these proteins. In particular, the protein OapC was previously hypothesized to bind a kink turn between J4a/6 and J5/6, and binding of OapC was thought to alter secondary structure, in particular increasing protection of J7/8 to in-line hydrolysis^17^. Our OLE dimer structure supports formation of a kink-turn in J4a/6 at the base of P5, but this kink turn is formed with J7/8, not J5/6 (**Fig. 1G**). The previously observed protection of J7/8 can therefore be explained by direct binding to the protein, and not by a rearrangement of secondary structure. In addition, whereas the internal loop of the P6 stem is different from the previously proposed one, it exposes residues 163-165, which were proposed to bind the protein RpsU^15^. A163 is flipped out of the helix and docks into a pocket created by P5, P6, P7, and dimer interface. This OLE dimer pocket is reminiscent of the pocket RpsU occupies in the ribosome, supporting the previous hypothesis that OLE could sequester RpsU^15^.

## ROOL assembles into an atomically ordered nanocage

ROOL is a class of RNAs encoded in a wide variety of bacterial prophages and phages, often near tRNA islands^6,11,12^. The predicted secondary structure is highly complex with multiple pseudoknots, but no protein binding partners have been identified, leading to the hypothesis that ROOL may function as an RNA-only complex^6^. Although no function has been described for ROOL, it has been shown to be as abundant as 16S ribosomal RNA, but non-essential, in at least one strain of *Ligilactobacillus salivarius*^12^.

The 659-nt ROOL *env-120*, discovered in cow rumen^6,27^, produces visually clear, symmetric particles in cryo-EM micrographs (**Fig. 2A, Extended Data** Fig. 3). The 3.1 Å reconstructed map reveals a closed, hollow nanocage structure comprising eight chains with dihedral symmetry and a diameter of approximately 280 Å, larger than the maximal dimension of a bacterial ribosome (∼250 Å) (**Fig. 2B** and **Supplemental Movie 2**). Each chain has a secondary structure consistent with the stems P1 to P19 proposed previously by covariation analysis^12^, including the pseudoknot P10 (**Fig. 2C**). Atomic models for each chain can be built with a good match with the map density as shown by the Q-scores^24^ (**Extended Fig. 3E**). Our model shows intramolecular tertiary interactions (**Fig. 2D-I**) which scaffold the flat monomer structure (**Fig. 2F-I**), including a set of non-canonical base pairs and stacking interactions connecting loops L3a and J6/7 (**Fig. 2F**), an A-minor interaction between L3c and P5 (**Fig. 2G**), an additional pseudoknot P13 (adjacent to P5 and P10, **Fig. 2H**), and a complex set of noncanonical pairs between nucleotides that are already in stems P1, P2, and P3b (**Fig. 2I**).

**Figure 2:**
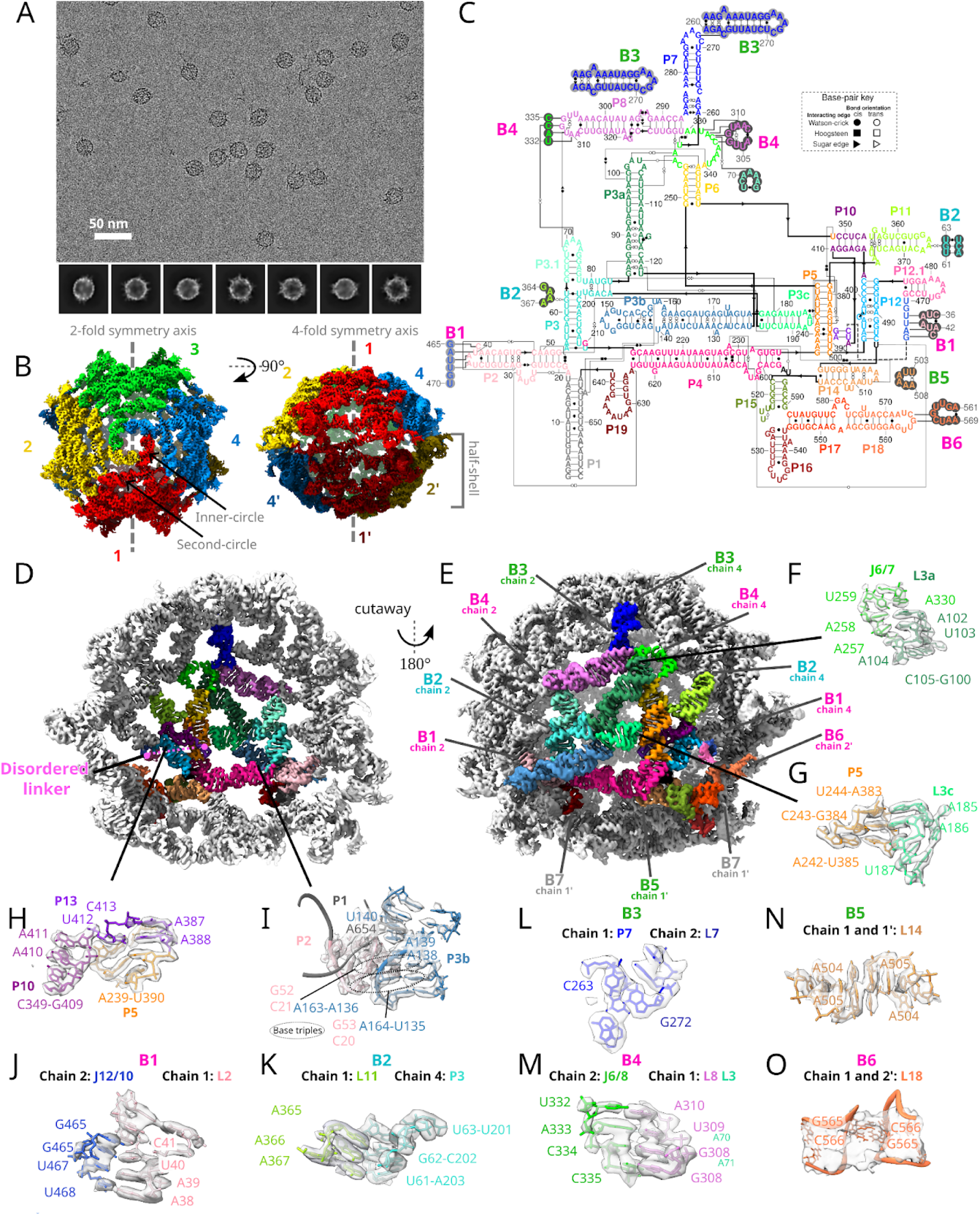
Atomically ordered structure of ROOL homo-octamer. (**A**) Representative micrograph with 2D class averages. (**B**) 3.1 Å cryo-EM reconstruction of the ROOL complex with D4 symmetry. The map is colored by 8 labeled chains. From the top-view, the inner and second circle are labeled. (**C**) Secondary structure of ROOL colored by domain. Only one chain is shown, in full. Nucleotides involved in intermolecular interactions have been circled in light or dark gray. (**D-E**) Chain 1 is colored by domain with all other chains in gray. (**D**) Cutaway to see this chain from the interior of the nanocage with the disordered linker labeled in pink. (**E**) Intermolecular interactions or bridges of Chain 1 are labeled, with kissing loops labeled in magenta, A-minor interactions in cyan, and other interactions in lime. One interaction, B7 (gray), is not ordered in this cryo-EM map, but residues come in close enough contact that interactions could form. The same interaction, but between different pairs of chains, share the same number. (**F-I**) Show selected intramolecular interactions. (**J-O**) Intermolecular interactions as labeled in (E).

**Figure 3:**
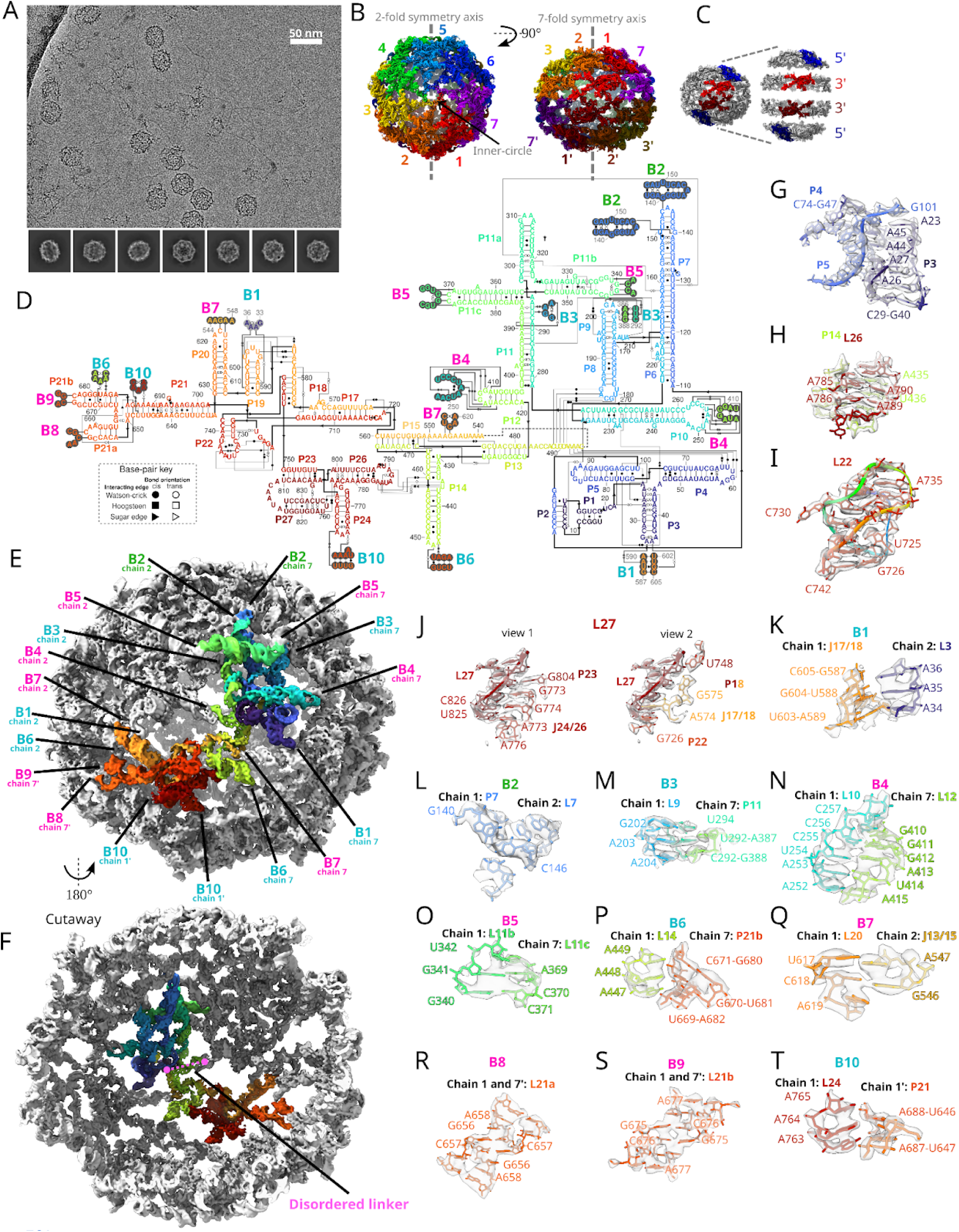
Atomically ordered structure of GOLLD homo-14-mer. (**A**) Representative micrograph and 2D class averages. (**B**) 3.0 Å cryo-EM reconstruction of GOLLD with D7 symmetry. The map is colored by chain. In the top-view, the inner circle is labeled. (**C**) The 5′ (blue, residues 1-420) and 3′ (red, residues 421-833) regions of GOLLD organize into the cap and a ring of the half-shell of the nanocage respectively. To demonstrate the separation of domains, the four regions are artificially moved apart. (**D**) The secondary structure of GOLLD. In the secondary structure diagram only one chain is displayed in full, nucleotides participating in intermolecular interactions that are from other chains are circled in light or dark gray. (**E-F**) One chain of GOLLD is colored rainbow, intermolecular interactions, or bridges, are labeled, with kissing loops labeled in magenta, A-minor interactions in cyan, and other interactions in lime. The same interaction, but between different pairs of chains, share the same number. Each chain interacts with four other chains. (F) is cutaway to see this chain from the interior of the nanocage with the disordered linker labeled in pink. (**G-J**) Select intramolecular interactions. (**K-T**) Intermolecular interactions as labeled in (F).

The ROOL quaternary complex is an octameric nanocage, with a top and bottom half-shells, each formed by four chains, hereafter labeled chains 1-4 and 1′-4′. Within a half-shell, each chain forms eight bridges with its neighbors, four on each side, labeled B1-B4 (**Fig 2J-O**). Starting from the top, the loop of stem P7 forms an isolated base-pair with a bulged out base in the next chain’s stem 7 (B3, **Fig. 2L**). This daisy chain of interacting stem-loops forms an inner circle on the top of the half-shell ∼36 Å in diameter (**Fig. 2B**). The P7 stem is not always conserved, but a second circle of RNA (**Fig. 2B**), involving a quaternary kissing loop (B4, **Fig. 2M**), is highly conserved in evolution as it was identified as tertiary interaction by previous covariation analysis^6^. An A-minor interaction (B2, **Fig. 2K**) and a novel quaternary kissing loop (B1, **Fig. 2J**) further glue together the chains in the half-shell. Between a novel intramolecular tertiary interaction (**Fig. 2G**) and the intermolecular kissing loop B1 (**Fig. 2J**), we identified a disordered region that appears to be located inside the nanocage, based on the position of flanking regions (**Figs. 2D**). This region was previously identified as a linker with little to no sequence or structural similarity across homologs^6^.

As opposed to a simple dimer like the OLE interface, where each chain interacts with a single partner, in the ROOL complex, each chain reaches over and interacts with two chains in the other half-shell. These interactions favor the full cage assembly, as opposed to isolated dimers. B5 and B6 are quaternary interactions where the same sequences but from different chains interact via adenosine stacking and Watson-Crick-Franklin base pairing, respectively (**Fig 2N-O**). An additional interaction between the internal loop J17/18 of chain 1, previously proposed to form a pseudoknot with the flank of the linker region, and P19 of chain 1′ seems plausible given their proximity, but that region was not resolved well in our structure.

## GOLLD assembles into a distinct atomically ordered nanocage

GOLLD RNAs are the largest among the three RNA classes analyzed here, with many members exceeding 800 nucleotides in length^8,11,28^. GOLLD, like ROOL, is a molecule of unknown function encoded in bacterial prophages and phages, often near tRNA islands, but with sequences and secondary structures distinct from those of ROOL^8,11,28^. GOLLD expression has been shown to increase during the lysis of bacterial cells infected by phage^11^. Unlike ROOL, the predicted secondary structures of GOLLD RNAs consist of a universally conserved 3′ region and a less conserved 5′ region^8^.

The GOLLD *env-38* RNA, discovered in a marine metagenomic sample downstream of Met-tRNA^8,29^, produces visually striking flower-like particles in cryo-EM micrographs (**Fig. 3A, Extended Data** Figure 4). 3D reconstruction at 3.0 Å resolution shows that GOLLD forms a nanocage, similar to ROOL, but larger. The GOLLD structure is a closed 14-mer with D7 quaternary symmetry, with a diameter of 380 Å and a completely empty interior (**Fig. 3B** and **Supplemental Movie 3**). Models for each of the 14 chains are built with Q-score^24^ in accordance with the map resolution (**Extended Data** Figure 4G). As with ROOL, kissing loops and A-minor interactions underlie the tertiary and quaternary structure of GOLLD (**Fig. 3D-T**), but the specific interactions are distinct. Beyond confirming the accuracy of the previously predicted secondary structure with stems P1-P27 (**Fig. 3D**), the tertiary structure of GOLLD reveals prominent interactions that have not been previously predicted, including A-minor interactions involving adenosines at the P3-P4-P5 junction (**Fig. 3G**), an A-minor interaction between adenosines in L26 and stem P14 (**Fig. 3H**), a loop L22 that forms a pseudoknot with the nearby linker J17/22, in addition to an A-minor interaction with that pseudoknot (**Fig. 3I**). Furthermore, loop L27 brings together 7 regions by forming base pairs with stem P23 and linker J24/26 as well as base-backbone interactions with two additional stems, P18 and P22, and linker J17/18 (**Fig. 3J**). Like in ROOL, the variable linker within each chain is not resolved, but the positions of immediate flanking sequences in the 5′ and 3′ regions indicate that the linker resides in the interior of the cage (**Fig. 3F**). Globally, the cryo-EM structure shows that the 5′ region and the 3′ region form separate domains in the 3D structure (**Fig. 3C**). This separation could explain why the 3′ and 5′ domains are divergent in GOLLD, whereas, in ROOL, the 5′ and 3′ regions are intertwined and hence have to co-evolve to maintain the tertiary and quaternary structure.

The 5′ domains of GOLLD form the cap of each half-shell of the nanocage. Within each half-shell’s cap, each of seven monomers forms eight quaternary bridges to other chains, 4 on each side, including kissing loops, A-minor interactions, and other interactions (B2-B5, **Fig. 3E,L-O**). B2 (**Fig. 3L**) is closely similar to the “daisy chain” of interacting stem-loops from ROOL, except that the distance between the interacting residues is reduced from 9 nt to 4 nt. This compensates for the increased number of chains in GOLLD, resulting in an inner circle of roughly the same diameter of ROOL. In GOLLD, the only non-interacting loop with a conserved sequence, L11a (sequence GAAA), points towards this inner circle. The 3′ regions complete the half-shell below this 5′ cap, through two interactions: an A-minor interaction (B6, **Fig. 3P**) and a kissing loop between L20 and J13/15 which was previously identified by covariation analysis^8^ and here shown to be an intermolecular bridge (B7, **Fig. 3Q**). Only a single intermolecular A-minor interaction, B1, glues the 3′ and 5′ regions from different chains together (**Fig. 3K**).

Lastly, similar to the ROOL nanocage, the two half-shells come together with each chain in the top half-shell interacting with two chains in the bottom half-shell. In the GOLLD nanocage, these interactions consist of two self-interacting kissing-loop interactions (B8 and B9, **Fig. 3R-S**) and an A-minor interaction (B10, **Fig. 3T**) involving 3′ regions from different chains.

## Biological relevance of homo-multimerization

Symmetric multimers are common among proteins and rationally designed RNA molecules^30^, but observations of natural RNA multimers are rare. When observed, natural RNA homomeric interactions typically involve a single contact^13^. Further, with notable exceptions of viruses, such as HIV and other retroviruses ^31^ and the Φ29 bacteriophage^32–34^, the biological relevance of RNA homomeric complexes has not been demonstrated, leaving the possibility that they form only at high RNA concentrations and extreme ionic conditions or in the context of the specific constructs chosen for *in vitro* structural characterization. In contrast, several lines of evidence support GOLLD, ROOL, and OLE forming multimers in their biological contexts.

First, mass photometry, which gives high precision estimates of molecular weight but requires molecular binding to surfaces, confirms the stoichiometry of GOLLD, ROOL, and OLE to be 14, 8, and 2, respectively, at RNA concentrations as low as 12.5 nM (**Extended Data** Fig. 6). This concentration is three orders of magnitude lower than the concentrations in our cryo-EM experiments; a population of only ∼10 molecules in a bacterial cell, substantially lower than what is expected from observed expression levels^12,16^.

Second, using dynamic light scattering (DLS; **Extended Data** Fig. 6), we confirmed that both ROOL and GOLLD primarily form thermostable multimers, with no detectable fraction of monomers, at temperatures up to 55 °C and concentrations as low as 110 nM.

Third, all three structures show five or more conserved inter-subunit contacts of each chain with other chains, complex arrangements that imply selection pressure during the evolution of these RNAs.

Fourth, concomitant with the same set of cryo-EM studies presented above, we resolved a 2.9 Å resolution map of another large RNA molecule, the *raiA* motif from *Clostridium acetobutylicum*^23,35^, as a pure monomer (**Extended Data** Figure 7), refuting the possibility that any large RNA would form a multimer in our experimental conditions. (An independent group has also resolved raiA as a monomer^36^.)

Fifth, using comparative analysis of both sequences and secondary structures, we detected evolutionary conservation of structural elements and, in particular, the sites of intermolecular interactions supporting RNA homo-oligomerization (**Supplemental File S1-S3** and **Supplementary Table S1**). Comparative analysis of OLE, ROOL and GOLLD showed that, although the sequences of these RNAs are not highly conserved, all intramolecular stems exhibit extensive base pairing supported by covariation analysis, including stems whose loops are involved in intermolecular bridges (**Extended Data** Fig. 8A-C**, Extended Data Text 2, and Supplemental Table 1**). The A positions in the OLE non-canonical A-A base pair stem bridge B1 and the GOLLD A-minor interaction bridge B6 are highly conserved (**Fig. 1D,3P** and **Extended Data** Fig. 8D-E). The intermolecular base pairs between ROOL J6/8 and L8 (bridge B4, **Fig. 2M**) was detected as a prominent, conserved quaternary interaction in prior covariation analysis^6^ (**Extended Data** Fig. 8B and **Supplemental Table 1**). In other bridges, we observed several intermolecular symmetric kissing loops with pairs between the same sequences from two different chains (**Extended Data** Fig. 8F-H). Apparent covariance at immediately adjacent nucleotides in these loop sequences would support intermolecular pairs because intramolecular canonical base pairing at adjacent nucleotides is stereochemically precluded and instead must be due to cross-chain pairings. OLE L9.3 and GOLLD L21a were each found to covary in this manner, switching an internal tetranucleotide between palindromes GGCC to GAUC or AGCU and an internal dinucleotide between GC and CG, supporting bridges OLE B3 and GOLLD B8, respectively (**Extended Data** Fig. 8F-G). The other symmetric kissing loops in our structures, GOLLD L21b (bridge B9) and ROOL L18 (bridge B6), were highly conserved across the variants for which the loops could be confidently aligned (**Extended Data** Fig. 8G-H), precluding similar covariance analysis but consistent with the importance of the observed intermolecular interactions.

## Discussion

Taken together, our cryo-EM data, biophysical experiments, and evolutionary analyses show that GOLLD, ROOL, and OLE each form not only ornate secondary structures but also symmetric quaternary assemblies stabilized by numerous intermolecular contacts, a type of RNA-only complex that has not been previously observed in nature. OLE forms a dimer shaped like a bundle of pipes and exposes structured binding pockets for protein partners. After superimposing an OapA dimer to each P4a site (OapA is known to bind OLE in a 2:1 ratio^22^), we note the RNA could induce the formation of an OapA tetramer. OapA is a membrane protein, and the tetramer is reminiscent of the dsRNA transporter SID-1^38–40^ suggesting that it may be able to accommodate RNA elements, such as the 3′ region of OLE, which was not resolved here (**Fig. 4A**). In contrast to OLE and despite unrelated sequences and distinct secondary and tertiary structures, GOLLD and ROOL both form nanocages, suggesting that their function might involve encapsulating other molecules, analogous to proteinaceous microcompartments that are common in bacteria and archaea^41^. Although not large enough to enclose entire DNA genomes of their parent phages, these cages might contain macromolecules of significant size (**Fig. 4B-C**), such as phage-encoded tRNAs, which are sometimes present in the GOLLD linker region, bacterial ribosomes, which have been shown to bind GOLLD in pull-down assays^11^, metabolites, or stress response proteins. It remains to be determined whether nanocage formation is a common feature among large natural RNAs.

**Figure 4:**
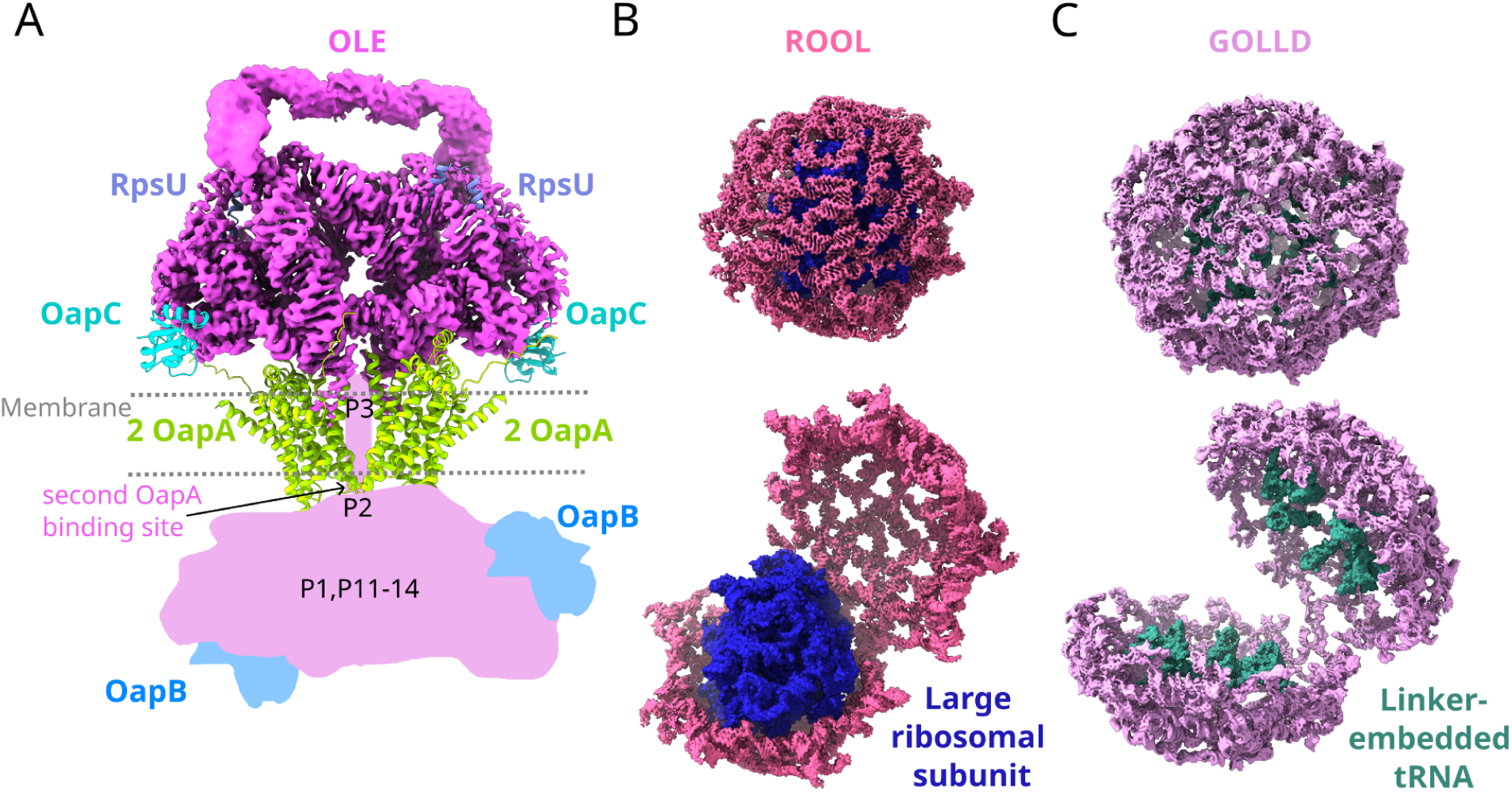
Structure-guided hypotheses for homo-oligomeric RNAs. (**A**) OLE dimer displayed with AlphaFold 3^37^ models of OapC and OapA dimer at their proposed binding sites. (**B-C**) The two half-shells of the RNA nanocages are held together by only a few interactions and hence could open up to encapsulate other biomolecules, such as protein, metabolites, nucleic acids or RNA-protein complexes. (**B**) The RNA nanocage formed by ROOL is shown encapsulating a ribosomal large subunit. (**C**) The RNA nanocages formed by GOLLD is shown exposing the covalently linked tRNAs when open.

## Acknowledgements

We thank Stanford Research Computing Center, SLAC Shared Scientific Data Facility, the Stanford-SLAC Cryo-EM Center, the Stanford Chem-H Macromolecular Structure Knowledge Center and the Stanford Biochemistry administrative staff for resources and support that contributed to this research. This work was supported by Stanford Bio-X (Bowes Graduate Student Fellowship to R.C.K.), the National Institute for Health (R35 GM122579 to R.D., Common Fund Transformative High-Resolution Cryo-Electron Microscopy program U24 GM129541 to W.C.), Howard Hughes Medical Institute (HHMI) (to R.D.), the National Science Foundation (Grant No. 2330652 to R.D. and W.C.), and the G. Harold & Leila Y. Mathers Foundation (MF-2303-04116 to A.G.). S.A.S. and E.V.K. report funding from the Intramural Research Program of the National Institutes of Health of the United States of America (National Library of Medicine). This article is subject to HHMI’s Open Access to Publications policy. HHMI lab heads have previously granted a nonexclusive CC BY 4.0 license to the public and a sublicensable license to HHMI in their research articles. Pursuant to those licenses, the author-accepted manuscript of this article can be made freely available under a CC BY 4.0 license immediately upon publication.

## Data availability

The cryo-EM micrographs and particles, cryo-EM maps, and model coordinates are being made available on EMPIAR, EMDB, and PDB, respectively. These data are available upon request until public release. Custom scripts can be found at https://github.com/DasLab/RNA_multimer_2024.

## Author contributions

R.C.K., W.C., and R.D. conceptualized and designed the study. R.C.K. and G.P.N. selected sequences for the study. V.W. and R.C.K. *in vitro* transcribed RNA and collected dynamic light scattering and mass photometry data. R.C.K. froze cryo-EM grids, collected and processed cryo-EM data, and modeled the cryo-EM maps with advice from W.C. and R.D.. G.P.N. screened cryo-EM grids. H.L. and A.G. generated sequence alignments and S.A.S., R.C.K., R.D., and E.V.K. analyzed these sequences for covariation. R.C.K., S.A.S., and R.D. prepared the manuscript with input from all authors.

## Methods

### *In vitro* RNA synthesis

DNA templates containing the RNA sequence of interest prepended with the T7 promoter (see **Extended Data Table 1** for sequence) were ordered as gBlocks from IDT. Primers designed to amplify this sequence (see **Extended Data Table 1** for sequence) were also ordered from IDT. PCR amplification was carried out with NEBNext Ultra II Q5 Master Mix (NEB #M0544S) using 10 ng of template per reaction. The thermocycler settings were: 98°C for 30 seconds, 35 cycles of 98°C for 10 seconds, 55°C for 30 seconds, 72°C for 30 seconds, and a final step of 72°C for 5 minutes. The PCR products were then column purified using the QIAquick PCR Purification Kit (Qiagen #28104) and run out on a 2% E-Gel agarose gel (Thermo Scientific #A42135) to check for DNA quality. DNA concentration was measured using a NanoDrop. Purified DNA smaller than 515 bp were in vitro transcribed using TranscriptAid T7 High Yield Transcription Kit (Thermo Scientific #K0441) with 6 μL of DNA template per reaction. Purified DNA longer than 515 bp were in vitro transcribed using MEGAscript T7 Transcription Kit (Thermo Scientific #AM1334) with 6 - 8 μL of DNA template per reaction. These in vitro transcription reactions were incubated at 37°C for 6 hours, then on 4°C hold before DNase treatment (respective of the transcription kit). The RNA was then purified using the RNA Clean & Concentrator-25 Kit (Zymo Research #R1017) and eluted in 30 μL of water. The concentration of purified RNA was measured using a NanoDrop, and the quality was checked using the Agilent 2100 Bioanalyzer (Nano RNA Assay, run by the Stanford PAN Facility), which has been plotted and displayed in Extended Data Figure 6A.

### RNA folding

For all subsequent experiments, RNA was re-folded using the same basic protocol. RNA concentrations used and any other modifications to this standard protocol are mentioned in each section. RNA was denatured (90 °C for 3 min, room temperature for 10 min) in 50 mM Na-HEPES pH 8.0. RNA was then folded with 10 mM MgCl_2_ at 50 °C for 20 min, and cooled to room temperature for at least 10 min before taking measurements.

### Mass photometry

Mass photometry data was collected using the Refeyn TwoMP, using AcquireMP version 2024-R1.1 and DiscoverMP version 2024-R1 to obtain histogram data. For the OLE data, coated glass slides from the MP Sample Preparation Pack (MP-CON-21014) were used, for the ROOL and GOLLD data the “Mass Glass UC” slides (MP-CON-41001) were used after coating with poly-L-lysine. The instrument was focused using droplet-dilution. Data were collected for 1 minute using the large image size. The contrast data were calibrated to nucleotide length using the Millennium RNA Markers (Thermo Scientific #AM7150). Gaussians were fitted by the automatic analysis in DiscoverMP. The resulting data and plotting code can be found in the accompanying GitHub repository.

Mass photometry data were not reliable for the *raiA* motif. raiA was folded at 1 µM following the standard procedure above. On the stage two dilutions were attempted, 15 µL buffer:2 µL sample for final concentration of 118 nM and 18 µL buffer:2 µL 10x diluted sample for a final concentration of 10 nM. There is a known issue with nucleic acid samples, whereby there are noisy low-mass peaks^42^ (communication with Refyn). These are not present in the buffer alone.

For this reason, *raiA* motif (205 nt) is below the recommended minimal size for mass photometry, and indeed when we attempted to collect data on the *raiA* motif we observed noise peaks, containing the size of raiA monomer but smaller than any multimer, in both binding and unbinding regimes, indicating unreliable results (data not shown).

OLE was folded at 0.25 µM and was diluted to 12.5 nM on the stage. OLE was folding in various buffers, all with 50 mM Na-HEPES pH 8.0 but various other components all added when MgCl_2_ is added in the standard protocol: (1) nothing added (2) 1 mM MgCl_2_ (3) 10 mM MgCl_2_ - standard (4) 100 mM MgCl_2_ (5) 10 mM MgCl_2_, 1% ethanol (6) 10 mM MgCl_2_, 5% ethanol (7) 0 mM MgCl_2_, 200 mM KCl (8) 10 mM MgCl_2_, 200 mM KCl (9) 0 mM MgCl_2_, 200 mM NaCl (10) 10 mM MgCl_2_, 200 mM NaCl. Buffers with MnCl_2_ were attempted but the manganese saturated the detector.

ROOL and GOLLD were folded at 1 µM. The samples were diluted 10x prior to taking data. On the stage the samples were further diluted (10 µL buffer: 10 µL sample) for a final concentration of 50 nM.

### Dynamic Light Scattering of RNA Nanocages

RNA was folded at 30 ng/µL using the standard folding protocol. Dynamic light scattering traces were collected using the Prometheus Panta. Two replicates (2 capillary of 10 µL volume, NanoTemper #PR-C002) for each RNA were obtained. Dynamic light scattering data of ten 5 second acquisitions per capillary with laser power 100% were obtained using PR.PantaControl v1.8.0. The auto-correlation function was calculated and size distribution was fitted using default parameters in PR.PantaAnalysis v1.8.0. The resulting size distribution tables and plotting code can be found in the accompanying GitHub repository.

### Cryo-electron Microscopy Grid Preparation

For all samples, the RNA was frozen using a VitroBot Mark IV, using #542 filter paper and Quantifoil 1.2/1.3 200 mesh copper grids which were glow discharged for 30 s at 15 mA. GOLLD was folded at 8 µM, using the standard folding conditions except, after the 50 °C incubation, the temperature was lowered to 37 °C at a rate of 0.1 °C/s, held at 37 °C for 2 min, and then reduced to room temperature at a rate of 0.1 °C/s. To increase concentration of GOLLD in the ice, 4 cycles of applying 2 µL of sample and blotting for 3 s were performed before plunging. ROOL was folded at 9.1 µM with the standard folding protocol. The grid was coated with 2 µL of 100 mM NaCl which was blotted for 3 s. Then, 2 µL sample was immediately applied to the grid and blotted for 3 s before plunging into liquid ethane. OLE and raiA RNA were frozen with the standard folding protocol at 20 µM and 15 µM respectively; 2 µL of sample was applied to the grid, followed by 3 s blot and plunge into liquid ethane.

### Cryo-electron Microscopy Data Collection

All datasets were collected on Titan Krios G3 microscopes using a 50 μm C2 aperture and 100 μm objective aperture and EPU software. The OLE dataset was collected using a Falcon 4 camera with a 10 eV slit on a Selectris energy filter, while the other datasets were collected using a K3 camera with a 20 eV slit on a Bio Quantum energy filter and EPU software. Additional information on dose, magnification, and data collected for each RNA can be found in **Extended Data Table 2**.

### Cryo-electron Microscopy Data Processing

Data were processed live using CryoSparc (v4.5.3)^43^ and then further refined, including non-uniform refinement^44^. For OLE and raiA per particle motion correction was performed^45^. For all datasets, symmetry was not applied until final refinement stages. For OLE, C2 symmetry was applied. For ROOL and GOLLD, D4 and D7 symmetry were applied respectively, followed by symmetry expansion of the particles and local refinement for one asymmetric unit. Finally, for GOLLD and ROOL subdomains of one asymmetric unit were locally refined and composite maps, and half-maps were created for one asymmetric unit and then composited to the full symmetry using phenix.combine_focused_maps (v1.21)^46^. Local resolution was estimated using CryoSparc. See **Extended Data** Figures 1-5 for more details on processing pipelines.

### Modeling

Maps were sharpened using phenix.auto_sharpen with half-maps (v1.21). Initial models for a monomer were obtained from ModelAngelo^47^; because current versions of ModelAngelo cannot be run on a pure RNA structure, EMDB-17659 was added to the corner of the map, the corresponding protein sequence (PDB: 8phe) was provided, and protein residues were subsequently deleted from the model. The RNA chains modeled were manually combined tracing the RNA sequences, adding and mutating residues when necessary (in particular, C to U mutations were commonly required). Manual model correction, and refinement was accomplished in Coot (version 0.9.8)^48^. Manual refinement of the monomer was performed using Isolde and Coot^48^. Symmetry was applied to the model, from henceforth refinement was done asymmetrically due to limitations in refinement programs. Intermolecular interactions were analyzed by hand and correct using Isolde^49^ and Coot^48^. DRRAFTER^50^ (Rosetta 3.10 (2020.42)) was used to fill in low resolution areas. For symmetric kissing loops, these models were selected and fit into the map and refined more symmetrically by hand using Isolde^49^. Final refinement was first run through phenix.real_space_refine^51^ followed by piecewise corrections using ERRASER2^52^ (Rosetta 3.10 (2020.42)), followed by manual refinement in Coot^48^ and Isolde^49^ when necessary. Validation metrics were calculated using Phenix, including phenix.rna_validate^53–55^. ChimeraX (version 1.8)^56^ was used to calculate Q-score^24^ and for all visuals. For visualizing a hypothetical RNA-protein complex, AlphaFold3^37^ was used to predict 1) a OapA dimer with a OapC monomer 2) RpsU using the sequences in (**Extended Data Table 1**). The OapA dimer was fitted into the proposed RNA site manually. The OapC was close to its binding site, but clashed with RNA, OapC the relative position of OapA and OapC was manually changed. RpsU was also placed manually in its proposed binding site. C2 symmetry was then applied to visualize the full complex.

### Bioinformatic analysis

Bacterial genomes were downloaded from National Center for Biotechnology Information (NCBI) Genome database in February 2024 (https://ftp.ncbi.nlm.nih.gov/genomes/genbank/bacteria/). GenBank records for phage genomes were downloaded in March 2024 (https://millardlab.org/bacteriophage-genomics/phage-genomes-march-2024/). Sequence profiles of GOLLD, ROOL, and OLE were downloaded from the Rfam database (ftp.ebi.ac.uk/pub/databases/Rfam/) on March 2024. A custom sequence profile of raiA was built using the reported alignment^35^. To retrieve ncRNAs from genome sequences, cmsearch was conducted using sequence profiles with a cutoff value of 1E-5^57^. The overall procedure yielded the following numbers of nonredundant ncRNA sequences: 806, GOLLD; 1,596, ROOL; 8,585, OLE; 4,875, raiA.

The Infernal software^57^ (v1.1.2) was used to compare candidate RNA structures against Rfam models (cmscan), build and calibrate new covariance models (cmbuild, cmcalibrate) for separate clusters of RNAs, and perform structure-informed homology searches (cmsearch). Comparative analysis and multiple alignments for isolated RNA candidates were conducted using cmalign^57^, MUSCLE^58^, with pairwise comparisons refined using the OWEN program^59^. Evolutionary history was inferred via the Maximum Likelihood method with different models in MEGA X^60^. Evolutionary analysis of compensatory substitutions in isolated clusters was performed by DecipherSSC^61^.

RNAalifold^62^ applied to computationally fold multiple RNA alignments, and Afold/Hybrid^63,64^ were used to predict locally folded secondary structures or hybrid duplex elements within clusters. Covariation analysis was performed with R-scape (v1.2.3)^65^, which annotates multiple structural alignments of RNAs using statistically significant covariations (E-value < 0.05) as base-pairing constraints.

---

**Extended Data Figure 1:**
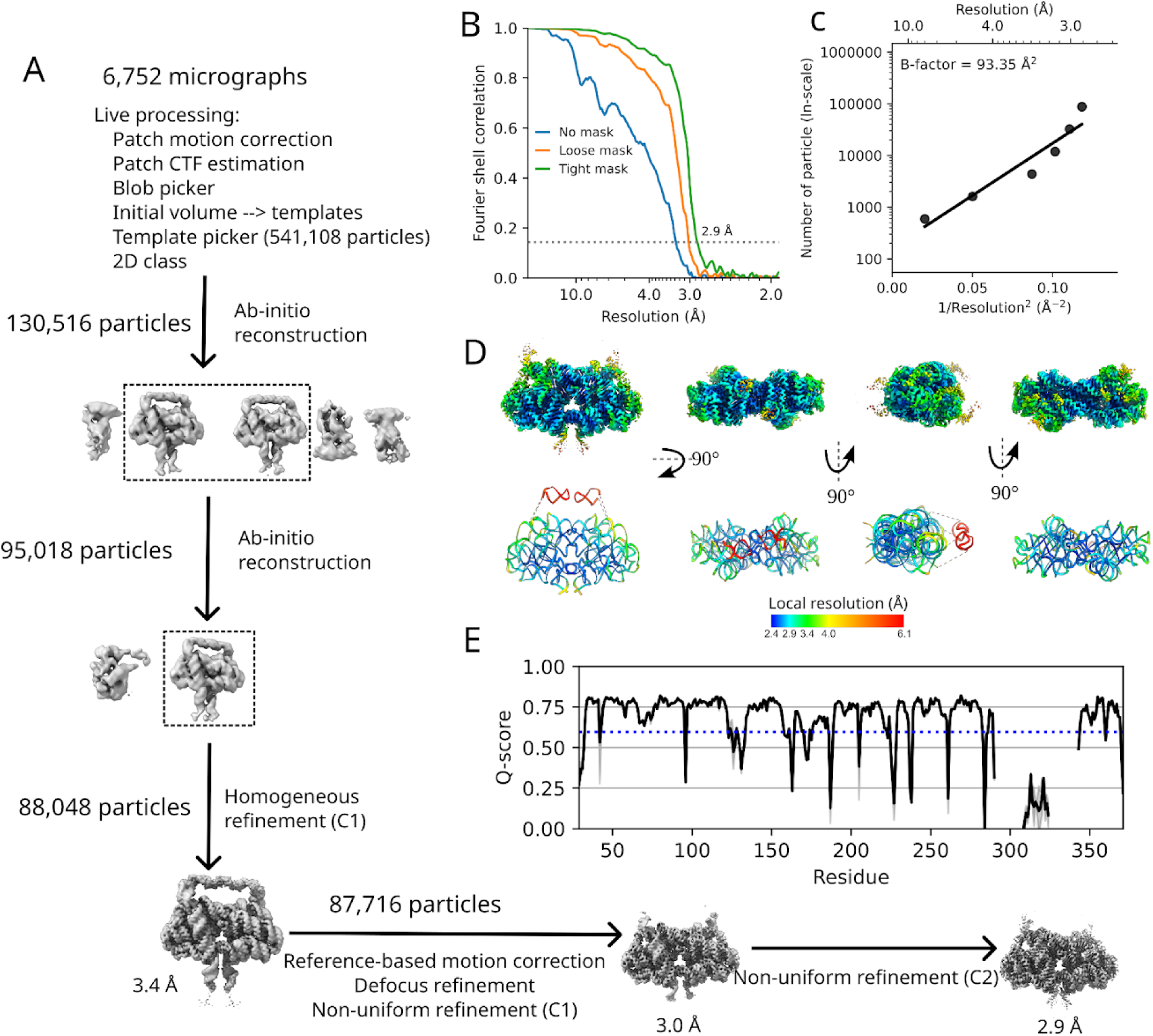
Cryo-EM data processing workflow for OLE dimer. (**A-E**) OLE resolves into a high resolution dimer, even in the absence of protein. (**A**) Data processing flowchart for the OLE dimer. (**B**) Fourier shell correlation (FSC) plot for final refinement of OLE dimer. (**C**) Plot of particle number against the reciprocal squared resolution for OLE dimer. The B-factor was calculated as twice the linearly fitted slope^1^. (**D**) Local resolution of the OLE dimer on the cryo-EM map (top) and the molecular model (bottom). (**E**) Resolvability of the built model of the OLE dimer as measured by Q-score. The black line is the mean across all chains, with the maximum and minimum values depicted by the shaded area. The expected Q-score at this resolution^2^ is labeled with a blue dotted line.

**Extended Data Figure 2:**
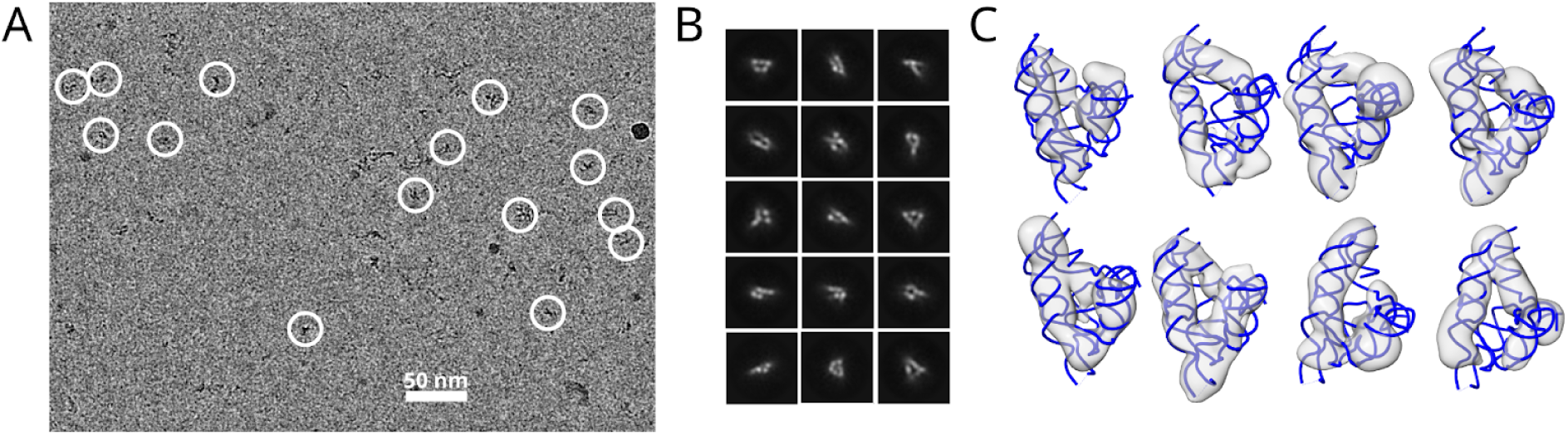
Cryo-EM data of HEARO RNA without protein. HEARO did not resolve into a high resolution structure, despite similar amount and quality of data as OLE-dimer. (**A**) The representative micrograph of HEARO shows clear particles. (**B**) Select 2D class averages show that HEARO is forming RNA helices, but they have diverse orientations and are blurred, suggesting high flexibility. (**C**) 3D reconstructions of HEARO, overlaid with the known structure of this RNA in the OMEGA nickase complex bound to protein IsrB (PDB: 8DMB^3^), show RNA of a similar fold to the complexed RNA. Multiple conformations are reconstructed, but with poorly resolved features, suggesting that HEARO may not form an atomically ordered structure when not in complex with its partner proteins.

**Extended Data Figure 3:**
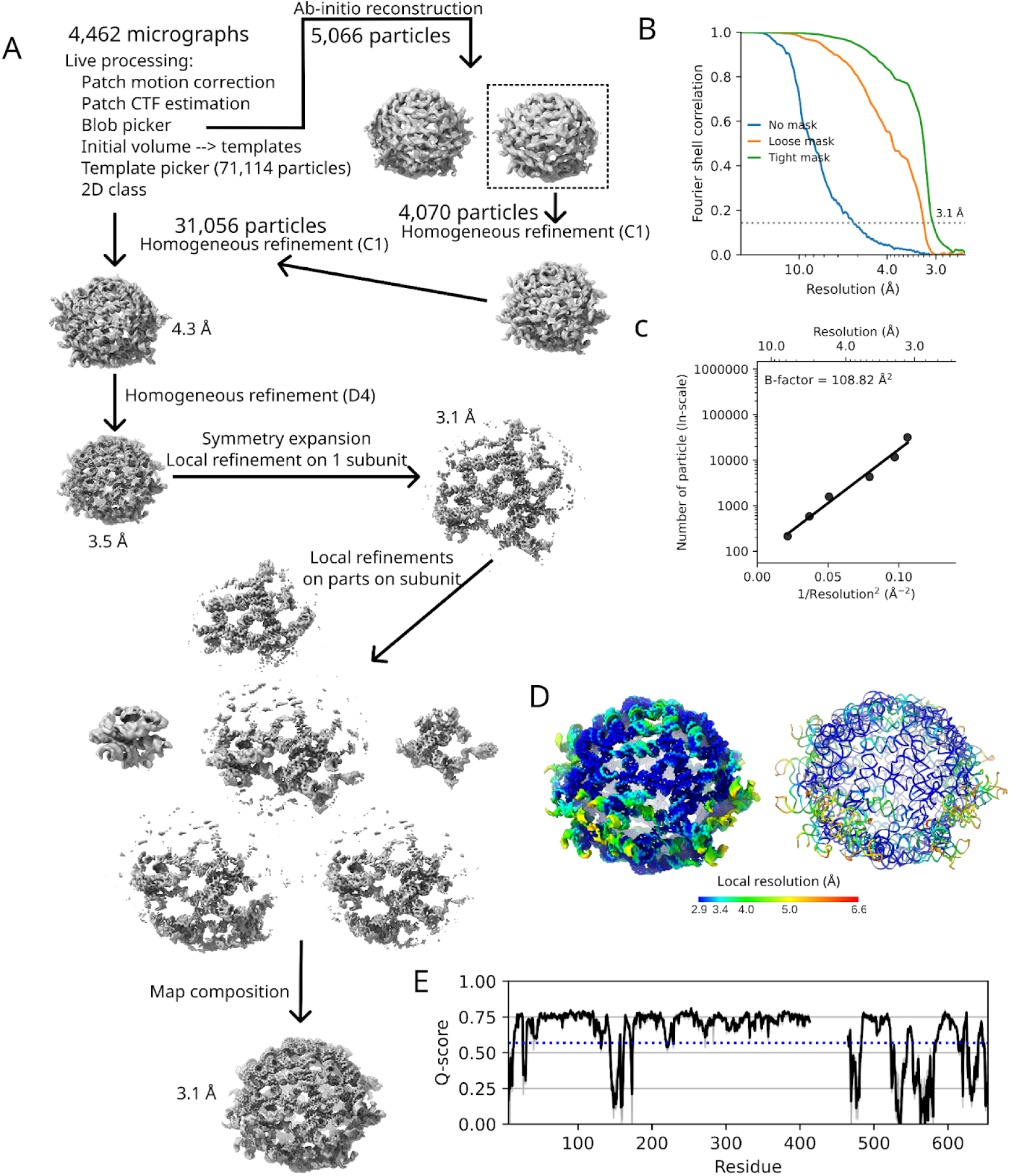
Cryo-EM data processing workflow for ROOL nanocage complex. (**A**) Data processing flowchart. (**B**) Fourier shell correlation (FSC) plots of the single subunit local refinement. (**C**) Plot of particle number against the reciprocal squared resolution for the single subunit local refinement. The B-factor was calculated as twice the linearly fitted slope^1^. (**D**) Local resolution on the cryo-EM map (right) and the molecular model (left). (**E**) Resolvability of the built model as measured by Q-score. The black line is the mean across all chains, with the maximum and minimum values depicted by the shaded area. The expected Q-score at this resolution^2^ is labeled with a blue dotted line.

**Extended Data Figure 4:**
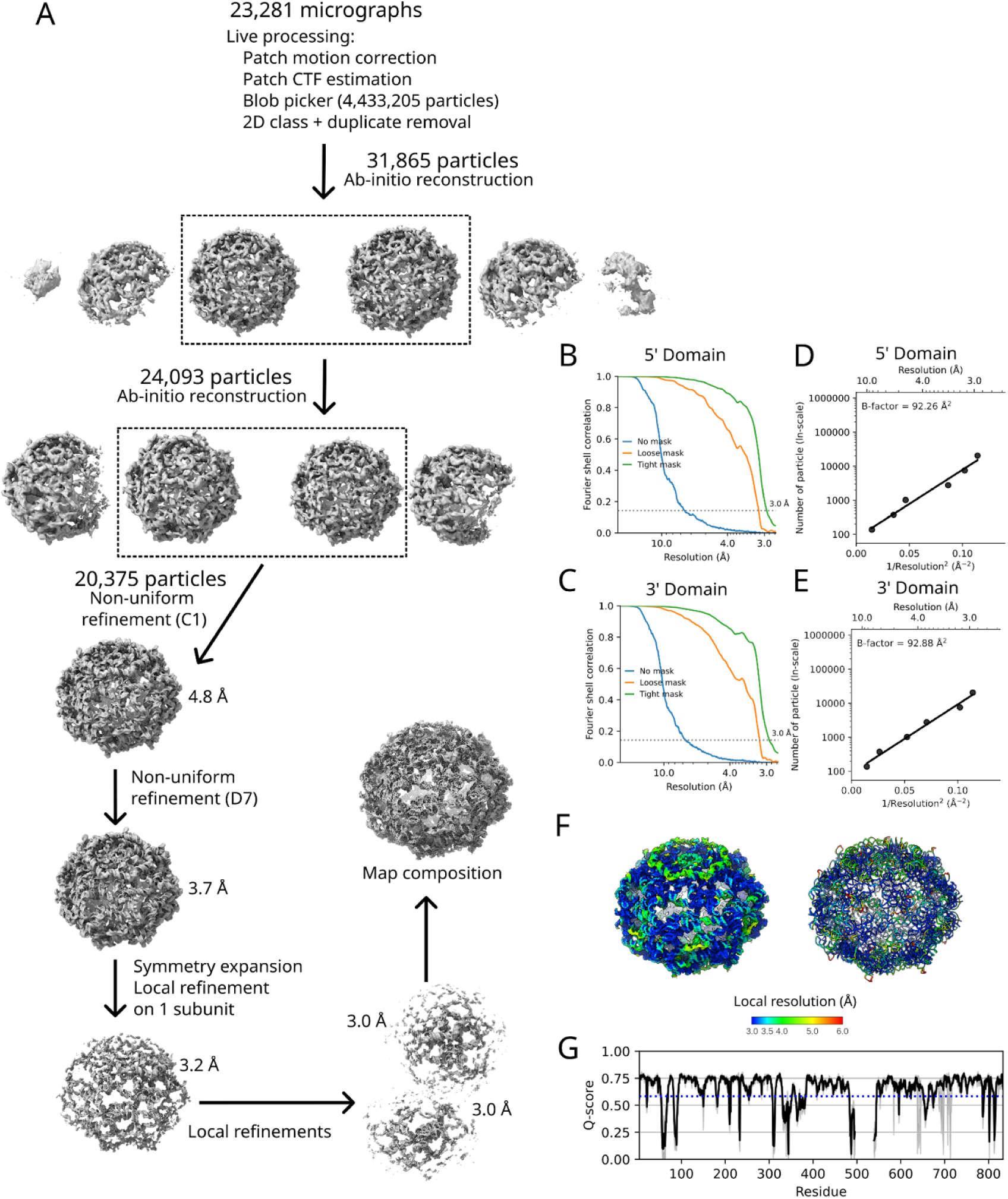
Cryo-EM data processing workflow for GOLLD nanocage complex. (**A**) Data processing flowchart. (**B**-**C**) Fourier shell correlation (FSC) plots for the local refinement of the 5′ and 3′ domains respectively. (**D-E**) Plots of particle number against the reciprocal squared resolution for the local refinement of the 5′ and 3′ domains respectively. The B-factor was calculated as twice the linearly fitted slope^1^. (**F**) Local resolution on the cryo-EM map (left) and the molecular model (right). (**G**) Resolvability of the built model as measured by Q-score. The black line is the mean across all chains, with the maximum and minimum values depicted by the shaded area. The expected Q-score at this resolution^2^ is labeled with a blue dotted line.

**Extended Data Figure 5:**
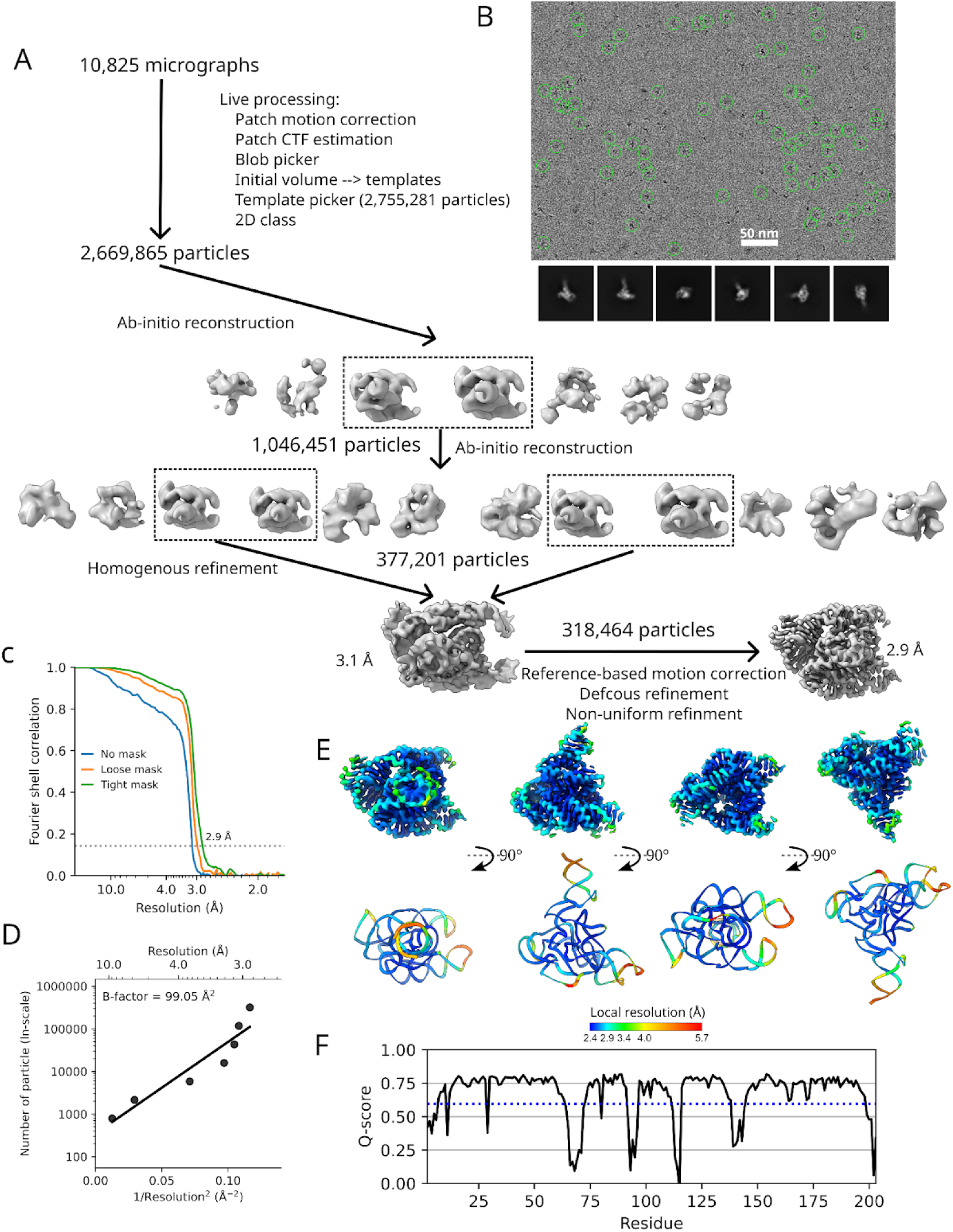
Cryo-EM data processing workflow for *raiA* motif. (**A**) Data processing flowchart. (**B**) Representative micrograph and 2D class averages. (**C**) Fourier shell correlation (FSC) plot. (**D**) Plot of particle number against the reciprocal squared resolution. The B-factor was calculated as twice the linearly fitted slope^1^. (**E**) Local resolution on the cryo-EM map (top) and the molecular model (bottom). (**F**) Resolvability of the built model as measured by Q-score. The expected Q-score of a RNA model at this resolution is labeled with a blue dotted line.

**Extended Data Figure 6:**
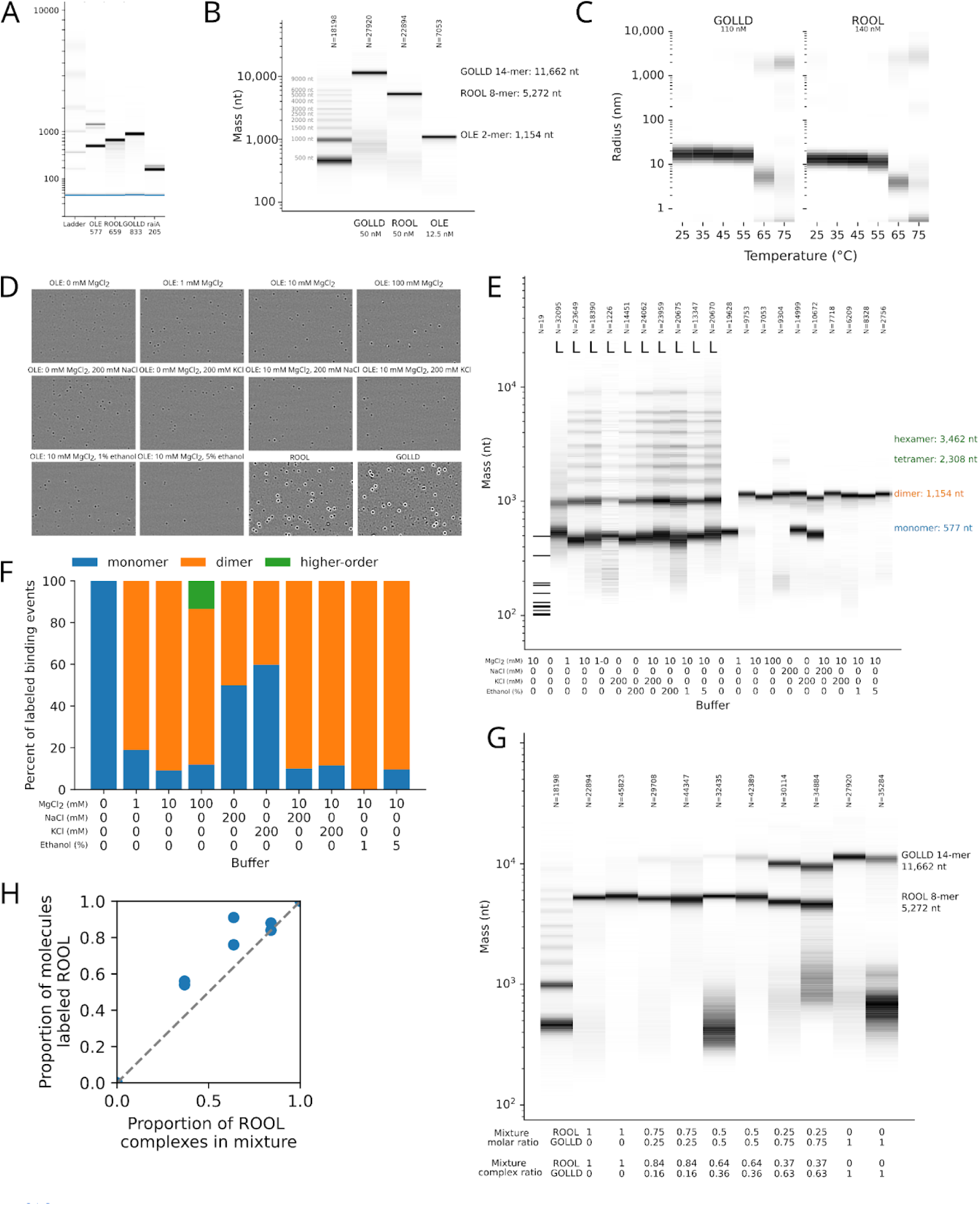
Evidence of multimer formation of GOLLD, ROOL, and OLE in biologically relevant concentrations. (**A**) Agilent Bioanalyzer traces demonstrate the purity of the samples. The second peak for OLE is a common artifact of poor denaturation of sample in Bioanalyzer traces. The pure monomeric reading in mass photometry, (B), shows that this peak is likely not a covalently linked dimer. (**B**) Mass of GOLLD, ROOL and OLE complexes as obtained from mass photometry at 50 nM, 50 nM and 12.5 nM respectively. The data is histogram of particle count density, normalized per sample, where dark is many counts, white is none. Total particle counts are shown above the graph. (**B**) Hydrodynamic radius of GOLLD and ROOL complexes as derived from dynamic light scattering at 110 nM and 140 nM respectively. The data is plotted as relative frequency, normalized by density per sample, with dark representing high frequency radii. The temperature of the sample was raised from 25 °C to 75 °C and dynamic light scattering traces were obtained every 10 °C, showing complex melting into monomers at 65 °C and aggregation at high temperatures. (**D**) Representative ratiometric image for all mass photometry data. (**E**) Mass photometry data of OLE in different buffer conditions demonstrates OLE can dimerize at low RNA concentration, low magnesium concentration, and in the absence of magnesium with sufficient monovalent cations. (**F**) The mass photometric data is summarized by counting the amount of hits in the monomer, dimer, and high stoichiometry peaks. The absolute ratio of monomer:dimer is accurate as assessed in (G-H). (**G**) Mass photometry traces of mixtures of ROOL and GOLLD, ratiometric image examples can be found in. (**H**) Summary of the mixture results, with the known complex ratio plotted against the ratio reported by mass photometry. A reasonable quantitative read-out is displayed, with slight bias towards higher counts for the smaller species, ROOL, opposite of the previously observed trend ^4^.

**Extended Data Figure 7:**
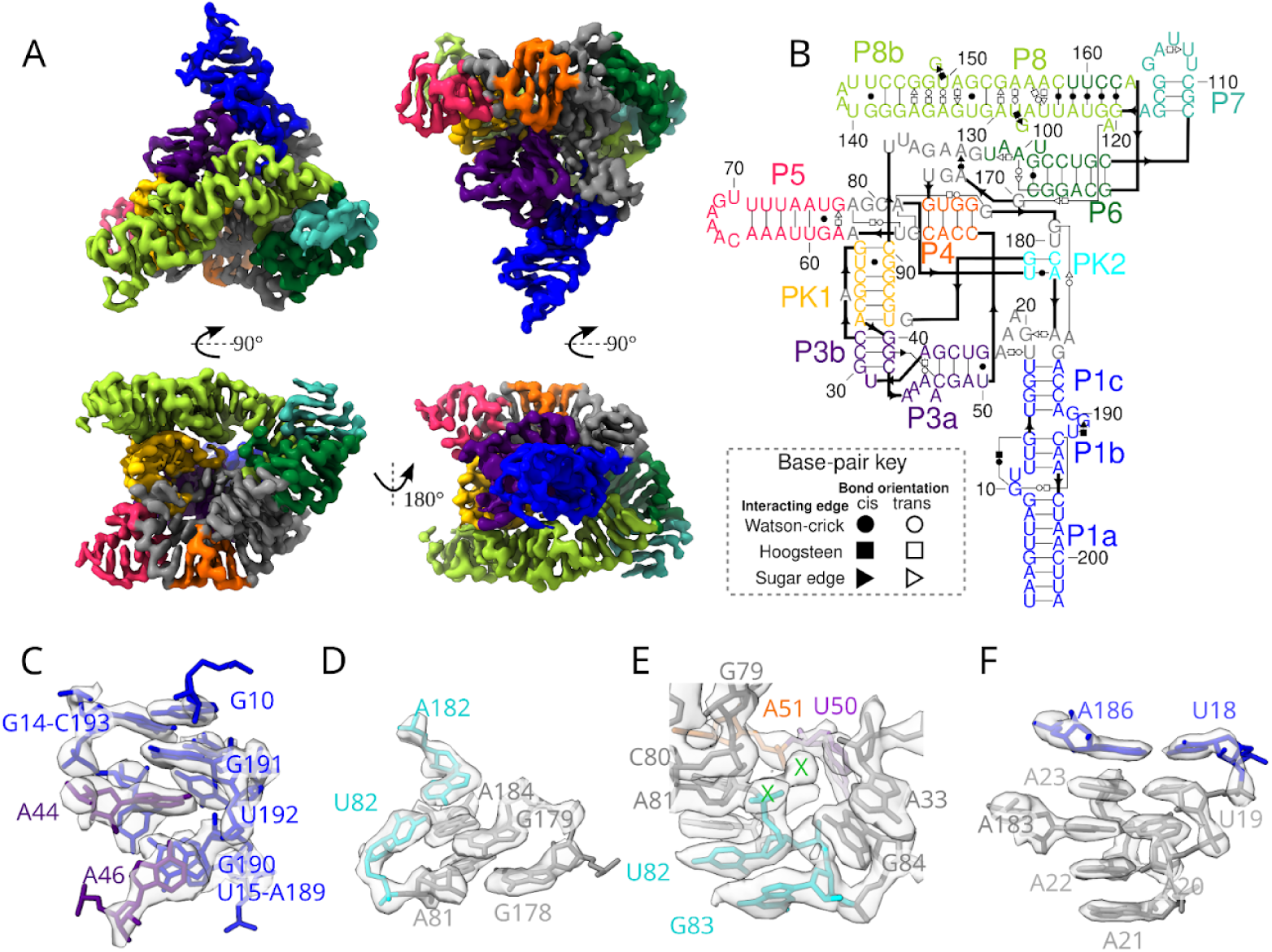
Tertiary structure of raiA RNA motif. (**A**) global view of tertiary structure of raiA motif and 2.9 Å cryo-EM map colored by as labeled in the secondary structure, (**B**). (**C-F**) Select tertiary interactions. Description can be found in **Extended Data Text 1**.

**Extended Data Figure 8:**
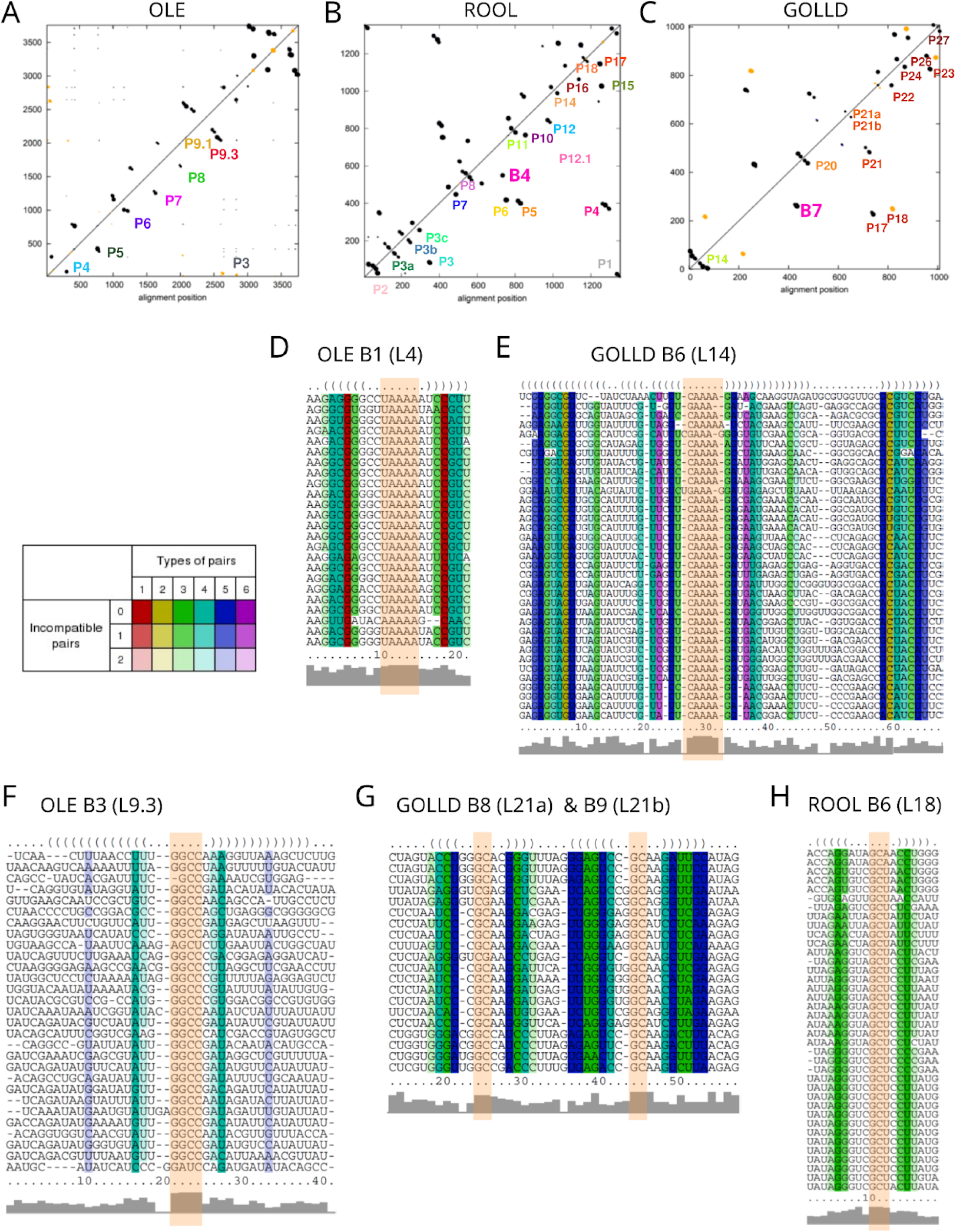
Comparative and covariation sequence analysis of homo-oligomer forming RNAs. (**A-C**) Distributions of covariation scores in multiple sequence alignments of (**A**) OLE, (**B**), ROOL, and (**C**) GOLLD sequences with select stems labeled. Dot size is proportional to the covariation score. In blue the consensus base pairs are depicted; in green, the consensus base pairs that show significant covariation are shown; in orange, other pairs that have significant covariation were depicted, they are not part of the consensus secondary structure but are compatible with it; in black, other significant pairs are depicted. Positions are relative to the original input alignment (before any gapped column is removed). (**D-H**) Examples of multiple alignments and profiles of sequence identity of selected stable hairpins with highly conserved loops which are involved in the intermolecular interactions are shown. Nucleotides involved in intermolecular interactions are labeled as in main Figures 1, 2 and 3 for the RNAs OLE (B1, B3), ROOL (B6) and GOLLD (B6, B8) respectively, and highlighted with an orange box. A coloring scheme for highlighting the mutational pattern with respect to the secondary structure (folding) was used and can be found next to (D). If one predicted base-pair is formed by several different combinations of nucleotides, consistent or compensatory mutations have taken place. This is indicated by different colors. Pale colors indicate that a base-pair cannot be formed in some sequences of the alignment. The sequence variants for the examples were selected from the closest branches of the evolutionary trees built based on the multiple sequence alignments used for the covariation analysis.

**Extended Data Table 1:**
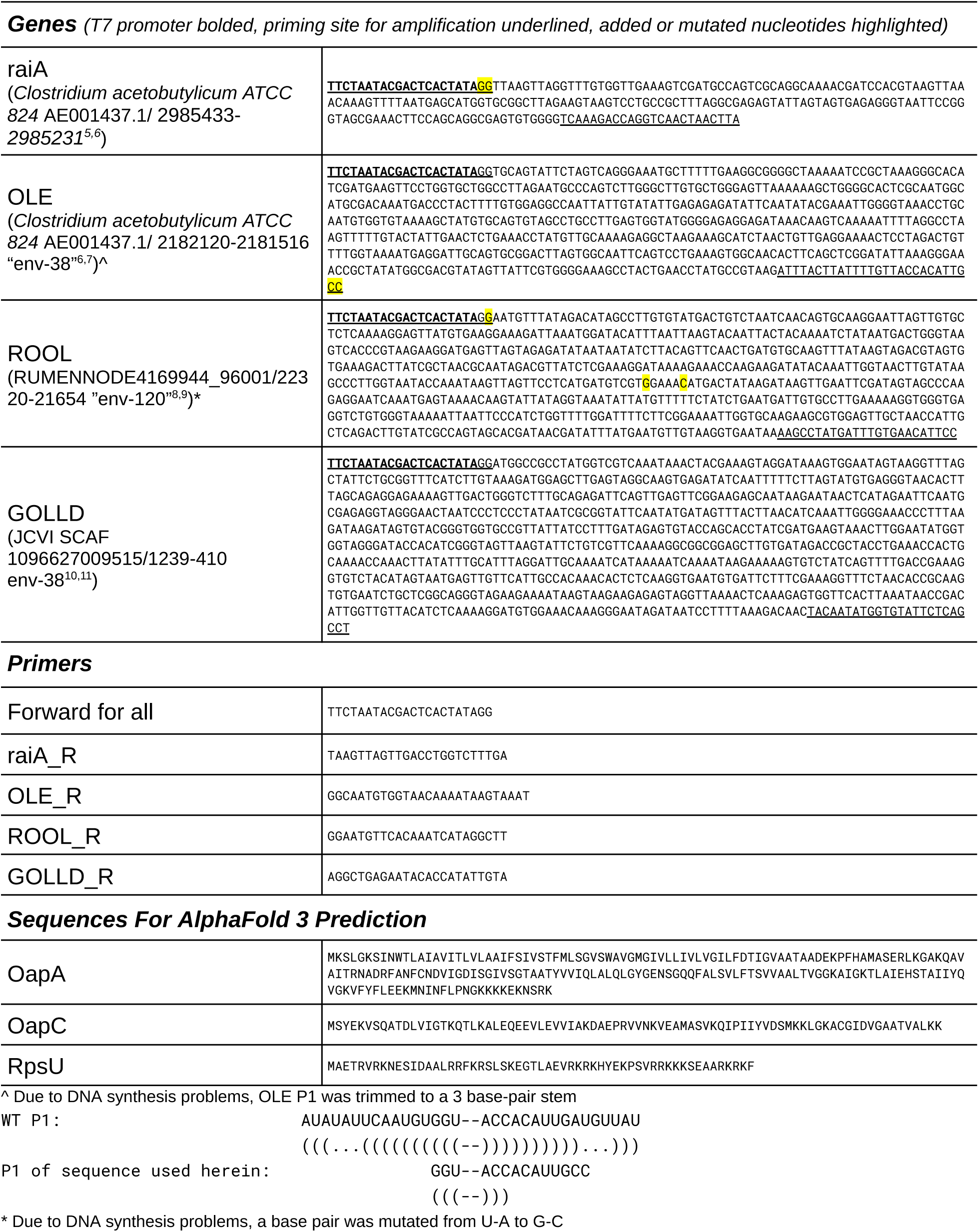
Sequences used in this study.

**Extended Data Table 2:** Cryo-EM experiments on four large non-coding RNAs.

|  | **OLE**<br>EMDB-48163<br>PDB-9MCW | **ROOL**<br>EMDB-48179<br>PDB-9MDS | **GOLLD**<br>EMDB-9MEE<br>PDB-48214 | ***raiA* motif**<br>EMDB-48162<br>PDB-9ELY |
| --- | --- | --- | --- | --- |
| **Sample preparation** |  |  |  |  |
| RNA length (nt) | 577 | 659 | 833 | 205 |
| Molecular weight (kDa) | 187 | 213 | 269 | 67 |
| Stoichiometry | 2 | 8 | 14 | 1 |
| Complex molecular weight (kDa) | 373 | 1,702 | 3,768 | 67 |
| Concentration (μM) | 15 | 9.1 | 8.0 | 20 |
| Concentration (mg/mL) | 2.8 | 1.9 | 2.1 | 1.3 |
| **Image acquisition** |  |  |  |  |
| Nominal magnification | 75,000 | 105,000 | 105,000 | 105,000 |
| Voltage (kV) | 300 | 300 | 300 | 300 |
| Electron exposure (e<sup>-</sup>/Å<sup>2</sup>) | 50 | 50 | 50 | 60 |
| Defocus range (μm) | -1.0 to -2.6 | -0.6 to -2.0 | -1.0 to -2.4 | -0.6 to -2.0 |
| Pixel size (Å) | 0.954 | 0.86 | 0.86 | 0.86 |
| Dose per frame (e<sup>-</sup>/Å<sup>2</sup>/frame) | 1.25 | 1.25 | 1.25 | 1.5 |
| Exposure time (s) | 4.74 | 1.42 | 2.00 | 1.65 |
| Micrographs acquired | 6,752 | 4,462 | 23,281 | 10,825 |
| **Data processing** |  |  |  |  |
| Symmetry imposed | C2 | D4 | D7 | C1 |
| Particle box size* (pix) (extraction size / fourier cropped size) | 360 / 240 | 600 / 400 | 800 / 500 | 320 / 320 |
| Initial particles images | 542,670 | 71,114 | 147,754 | 2,755,281 |
| Final particle images | 87,716 | 31,656 | 20,357 | 318,464 |
| Map resolution (Å) FSC threshold 0.143 | 2.9 | 3.1 | 3.0 | 2.9 |
| Map resolution range (Å) | 2.4-6.1 | 2.9-6.6 | 3.0-6.0 | 2.4-5.7 |
| B-factor (Å<sup>2</sup>) | 93 | 109 | 93 | 99 |
| **Modeling** |  |  |  |  |
| Non-hydrogen atoms | 13,206 | 102,240 | 235,872 | 4,318 |
| RNA nucleotides | 616 | 4,784 | 11,032 | 202 |
| Average Q-score | 0.66 | 0.63 | 0.66 | 0.68 |
| Model resolution (Å) FSC threshold 0.5 | 3.04 | 3.38 | 3.18 | 3.11 |
| Molprobity score | 1.73 | 1.98 | 2.13 | 1.71 |
| R.m.s.d. bond lengths (Å) | 0.015 | 0.009 | 0.010 | 0.003 |
| R.m.s.d. bond angles (°) | 1.583 | 1.193 | 1.184 | 0.703 |
| RNA sugar pucker outliers | 9 (1%) | 152 (3%) | 220 (2%) | 0 (0%) |
| RNA suite outliers | 41 (7%) | 621 (13%) | 1527 (14%) | 16 (8%) |
| RNA suiteness | 0.684 | 0.641 | 0.623 | 0.735 |
| Clashscore | 0.20 | 1.20 | 2.11 | 0.15 |

\*additional box sizes used during processing

**Extended Data Text 1:** Description of raiA RNA tertiary structure.

the structure of another family of long RNAs, the *raiA* motif, from *Clostridium acetobutylicum* ATCC 824, was determined. Under similar concentrations and conditions to the other long RNAs, the raiA motif was found to be a well-ordered monomer (**Extended Data** Fig. 7A). The secondary structure of the raiA RNA is very similar to the structure proposed by Soares et al^5^ based on covariance analysis and inline probing experiments (**Extended Data** Fig. 7B). The few differences are trivial, four previously proposed base pairs were not found in our structure: U19-G185, U29-A43, U50-G178, G55-U173, and an additional four base pairs were identified not previously annotated, mostly within the region for which there was no prior data from inline probing assays: G24-U50, G101-C169, C103-G168, G120-C162. Notably, although U29-A43 was proposed to be a base-pair, both bases were sites of spontaneous scission supporting our model with no base pair.

The linkers J1a/1b, J1b/1c and J3a/3b all interact in a complex tertiary interaction rigidifying P1 and P3 (**Extended Data** Figure 7C). Around PK2, U82-A182 and G83-C181, our structure shows an interesting interaction where the strand loops around itself to form non-canonical interaction A184-G179 which facilitates the sharp turn of the backbone necessary to form this pseudoknot (**Extended Data** Fig. 7D). The backbone of the 3′ strand of PK2 is part of an interesting pocket. The pocket contains two cryo-EM densities of high signal indicating ions (**Extended Data** Fig. 7E). These are 6 Å apart. The pocket is formed by backbone atoms as well as G84, which is observed pointing into the pocket.

There are two major secondary structure variants in the raiA motif family; both are consistent with our 3D structure. In ∼30% of raiA motif sequences, P7 and P8 are not present^1^. These stems appear peripheral, wrapping around the core of the RNA. There is weak density in the cryo-EM map indicating the P8-loop may interact with the P5-loop; however this region has poor local resolution, suggesting the interaction is transient or flexible and hence does not play an important stabilizing role for the atomically ordered core. Second, the loop in J1c/3a (19-23) forms a T-loop, into which A183 intercalates (**Extended Data** Fig. 7F). ∼40% of raiA motifs have an additional stem, P2, between P1c and P3a^1^, which would not support a T-loop. We hypothesize that the T-loop interaction is replaced by an A-minor interaction between the P2 minor groove and A183 and/or A184 (>97% sequence conserved^1^). Hence, we hypothesize that these secondary structure variants would likely have similar folds to the structure shown herein.

The long stretch of nucleotides, Jpk1/6, was previously identified as having many conserved nucleotides but being unpaired according to covariation and scission experiments^1^. We observe low local resolution in this region, supporting the lack of structure and hence flexibility of this region. Interestingly, this region is looped out into solution as opposed to being tucked into the core as might be expected for regions with high sequence conservation. Hence, this sequence does not seem to play a vital role in the structure of this RNA, and the sequence may be an important recognition signal for other nucleic acid or protein partners. Alternatively, it may play an important structural role in an alternative biologically important conformation of this molecule that may be visited at different stages of this RNA’s function, which remains unknown.

**Extended Data Text 2:** Comparative and covariation analysis of ornate large RNAs.

To gain insight into the biological significance and divergence of protein-independent RNA quaternary ensembles, we analyzed the evolutionary conservation of their intermolecular interactions across distinct families of large RNA molecules. This included comparative and covariance analyses of the intermolecular contacts identified in this study and the structural elements in their neighborhoods. The analysis of OLE, ROOL, and GOLLD showed that, despite limited sequence identity, they all exhibit extensive folding and some interactions that are long range in sequence, as confirmed by covariation statistics (**Extended Data** Fig. 8A-C and **Supplemental Files 1-3**). Covariance analysis across the three large RNA families revealed similar patterns in the distribution of covariance scores (**Extended Data** Fig. 8A-C and **Supplemental Table 1**), with significant enrichment of covariance scores at sites close to identified intermolecular interactions. Notably, most of the consensus positions lacked significant covariation scores due to the requirement of some variability to be able to observe sufficient mutations to detect covariation. Instead, pairs with significant covariation scores (*E*<0.05) tended to be located in variable regions of hairpin stems, while their loops remained highly conserved (**Supplemental Table 1**). This pattern underscores the role of covariation in maintaining structural stability despite sequence variability.

We observed that some hairpins with variable stems maintained highly conserved loops, particularly those containing A-runs, for example, B1 interaction in OLE (P4/L4) and B6 interaction (L14/P14) in GOLLD (see Fig. 3 and **Extended Data** Fig. 8D-E). In these examples, the four or three highly conserved A positions (**Extended Data** Fig. 8D-E), flanked by variable stems with a number of significant covariations (**Supplemental Table 1**), are indeed involved in A-minor interactions in our cryo-EM structures (B1 in Fig. 1D, B6 in Fig. 3P, respectively).

**Supplemental Table 1:** Summary of the nucleotide-nucleotide covariations identified in the OLE, ROOL, and GOLLD alignments which can be found in **Supplemental Files 1, 2, and 3** respectively.

**Supplemental File 1:** The multiple sequence alignment of OLE in stockholm format.

**Supplemental File 2:** The multiple sequence alignment of ROOL in stockholm format.

**Supplemental File 3:** The multiple sequence alignment of GOLLD in stockholm format.

**Supplemental Video 1:** The overall topology of the OLE dimer is displayed with the regions in Fig. 1 highlighted.

**Supplemental Video 2:** The overall topology of the ROOL nanocage is displayed with the regions in Fig. 2 highlighted.

**Supplemental Video 3:** The overall topology of the GOLLD nanocage is displayed with the regions in Fig. 3 highlighted.

## References

1. ENCODE Project Consortium. An integrated encyclopedia of DNA elements in the human genome. Nature 489, 57–74 (2012).

2. Nemeth, K., Bayraktar, R., Ferracin, M. & Calin, G. A. Non-coding RNAs in disease: from mechanisms to therapeutics. Nat. Rev. Genet. 25, 211–232 (2024).

3. Eddy, S. R. Non-coding RNA genes and the modern RNA world. Nat. Rev. Genet. 2, 919–929 (2001).

4. Stav, S. et al. Genome-wide discovery of structured noncoding RNAs in bacteria. BMC Microbiol. 19, 66 (2019).

5. Chen, Y. et al. Hovlinc is a recently evolved class of ribozyme found in human lncRNA. Nat. Chem. Biol. 17, 601–607 (2021).

6. Weinberg, Z. et al. Detection of 224 candidate structured RNAs by comparative analysis of specific subsets of intergenic regions. Nucleic Acids Res. 45, 10811–10823 (2017).

7. Puerta-Fernandez, E., Barrick, J. E., Roth, A. & Breaker, R. R. Identification of a large noncoding RNA in extremophilic eubacteria. Proc. Natl. Acad. Sci. U. S. A. 103, 19490–19495 (2006).

8. Weinberg, Z., Perreault, J., Meyer, M. M. & Breaker, R. R. Exceptional structured noncoding RNAs revealed by bacterial metagenome analysis. Nature 462, 656–659 (2009).

9. Narunsky, A. et al. The discovery of novel noncoding RNAs in 50 bacterial genomes. Nucleic Acids Res. 52, 5152–5165 (2024).

10. Ontiveros, N. et al. Rfam 15: RNA families database in 2025. Preprint bioRxiv 2024.09.23.614430 (2024).

11. Chen, A. G. Functional Investigation of Ribozymes and Ribozyme Candidates in Viruses, Bacteria and Eukaryotes. (Yale University, 2014).

12. Cousin, F. J. et al. A long and abundant non-coding RNA in Lactobacillus salivarius. Microb Genom 3, e000126 (2017).

13. Jones, C. P. & Ferré-D’Amaré, A. R. RNA quaternary structure and global symmetry. Trends Biochem. Sci. 40, 211–220 (2015).

14. Lyon, S. E., Wencker, F. D. R., Fernando, C. M., Harris, K. A. & Breaker, R. R. Disruption of the bacterial OLE RNP complex impairs growth on alternative carbon sources. PNAS Nexus 3, gae075 (2024).

15. Breaker, R. R., Harris, K. A., Lyon, S. E., Wencker, F. D. R. & Fernando, C. M. Evidence that OLE RNA is a component of a major stress-responsive ribonucleoprotein particle in extremophilic bacteria. Mol. Microbiol. 120, 324–340 (2023).

16. Wallace, J. G., Zhou, Z. & Breaker, R. R. OLE RNA protects extremophilic bacteria from alcohol toxicity. Nucleic Acids Res. 40, 6898–6907 (2012).

17. Lyon, S. E., Harris, K. A., Odzer, N. B., Wilkins, S. G. & Breaker, R. R. Ornate, large, extremophilic (OLE) RNA forms a kink turn necessary for OapC protein recognition and RNA function. J. Biol. Chem. 298, 102674 (2022).

18. Widner, D. L., Harris, K. A., Corey, L. & Breaker, R. R. Bacillus halodurans OapB forms a high-affinity complex with the P13 region of the noncoding RNA OLE. J. Biol. Chem. 295, 9326–9334 (2020).

19. Yang, Y., Harris, K. A., Widner, D. L. & Breaker, R. R. Structure of a bacterial OapB protein with its OLE RNA target gives insights into the architecture of the OLE ribonucleoprotein complex. Proc. Natl. Acad. Sci. U. S. A. 118, e2020393118 (2021).

20. 20. Fernando, C. M. & Breaker, R. R. Bioinformatic prediction of proteins relevant to functions of the bacterial OLE ribonucleoprotein complex. mSphere 9, e0015924 (2024).

21. Harris, K. A., Zhou, Z., Peters, M. L., Wilkins, S. G. & Breaker, R. R. A second RNA-binding protein is essential for ethanol tolerance provided by the bacterial OLE ribonucleoprotein complex. Proc. Natl. Acad. Sci. U. S. A. 115, E6319–E6328 (2018).

22. Block, K. F., Puerta-Fernandez, E., Wallace, J. G. & Breaker, R. R. Association of OLE RNA with bacterial membranes via an RNA-protein interaction. Mol. Microbiol. 79, 21–34 (2011).

23. Nölling, J. et al. Genome sequence and comparative analysis of the solvent-producing bacterium Clostridium acetobutylicum. J. Bacteriol. 183, 4823–4838 (2001).

24. Pintilie, G. et al. Measurement of atom resolvability in cryo-EM maps with Q-scores. Nat. Methods 17, 328–334 (2020).

25. Hirano, S. et al. Structure of the OMEGA nickase IsrB in complex with ωRNA and target DNA. Nature 610, 575–581 (2022).

26. Frank, J. et al. A model of the translational apparatus based on a three-dimensional reconstruction of the Escherichia coli ribosome. Biochem. Cell Biol. 73, 757–765 (1995).

27. Hess, M. et al. Metagenomic discovery of biomass-degrading genes and genomes from cow rumen. Science 331, 463–467 (2011).

28. Morgado, S., Antunes, D., Caffarena, E. & Vicente, A. C. The rare lncRNA GOLLD is widespread and structurally conserved among Mycobacterium tRNA arrays. RNA Biol. 17, 1001–1008 (2020).

29. Yooseph, S., et al. The Sorcerer II Global Ocean Sampling expedition: expanding the universe of protein families. PLoS Biol. 5, e16 (2007).

30. Hao, C. et al. Construction of RNA nanocages by re-engineering the packaging RNA of Phi29 bacteriophage. Nat. Commun. 5, 3890 (2014).

31. Dubois, N., Marquet, R., Paillart, J.-C. & Bernacchi, S. Retroviral RNA dimerization: From structure to functions. Front. Microbiol. 9, 527 (2018).

32. Ding, F. et al. Structure and assembly of the essential RNA ring component of a viral DNA packaging motor. Proc. Natl. Acad. Sci. U. S. A. 108, 7357–7362 (2011).

33. Simpson, A. A. et al. Structure of the bacteriophage phi29 DNA packaging motor. Nature 408, 745–750 (2000).

34. Xu, J., Wang, D., Gui, M. & Xiang, Y. Structural assembly of the tailed bacteriophage ϕ29. Nat. Commun. 10, 2366 (2019).

35. Soares, L. W., King, C. G., Fernando, C. M., Roth, A. & Breaker, R. R. Genetic disruption of the bacterial *raiA* motif noncoding RNA causes defects in sporulation and aggregation. Proc. Natl. Acad. Sci. U. S. A. 121, e2318008121 (2024).

36. Badepally, N. G., de Moura, T. R., Purta, E., Baulin, E. F. & Bujnicki, J. M. Cryo-EM structure of raiA ncRNA from Clostridium reveals a new RNA 3D fold. J. Mol. Biol. 436, 168833 (2024).

37. Abramson, J. et al. Accurate structure prediction of biomolecular interactions with AlphaFold 3. Nature 630, 493–500 (2024).

38. Zheng, L. et al. Cryo-EM structures of human SID-1 transmembrane family proteins and implications for their low-pH-dependent RNA transport activity. Cell Res. 34, 80–83 (2024).

39. Hirano, Y. et al. Cryo-EM analysis reveals human SID-1 transmembrane family member 1 dynamics underlying lipid hydrolytic activity. *Commun*. Biol. 7, 664 (2024).

40. Qian, D. et al. Structural insight into the human SID1 transmembrane family member 2 reveals its lipid hydrolytic activity. Nat. Commun. 14, 3568 (2023).

41. Greening, C. & Lithgow, T. Formation and function of bacterial organelles. Nat. Rev. Microbiol. 18, 677–689 (2020).

42. Li, Y., Struwe, W. B. & Kukura, P. Single molecule mass photometry of nucleic acids. Nucleic Acids Res. 48, e97 (2020).

43. Punjani, A., Rubinstein, J. L., Fleet, D. J. & Brubaker, M. A. cryoSPARC: algorithms for rapid unsupervised cryo-EM structure determination. Nat. Methods 14, 290–296 (2017).

44. Punjani, A., Zhang, H. & Fleet, D. J. Non-uniform refinement: adaptive regularization improves single-particle cryo-EM reconstruction. Nat. Methods 17, 1214–1221 (2020).

45. Rubinstein, J. L. & Brubaker, M. A. Alignment of cryo-EM movies of individual particles by optimization of image translations. J. Struct. Biol. 192, 188–195 (2015).

46. Liebschner, D. et al. Macromolecular structure determination using X-rays, neutrons and electrons: recent developments in Phenix. Acta Crystallogr. D Struct. Biol. 75, 861–877 (2019).

47. Jamali, K. et al. Automated model building and protein identification in cryo-EM maps. Nature 628, 450–457 (2024).

48. Emsley, P., Lohkamp, B., Scott, W. G. & Cowtan, K. Features and development of coot. Acta Crystallogr. D Biol. Crystallogr. 66, 486–501 (2010).

49. Croll, T. I. ISOLDE: a physically realistic environment for model building into low-resolution electron-density maps. Acta Crystallogr D Struct Biol 74, 519–530 (2018).

50. Kappel, K. et al. De novo computational RNA modeling into cryo-EM maps of large ribonucleoprotein complexes. Nat. Methods 15, 947–954 (2018).

51. Afonine, P. V. et al. Real-space refinement in PHENIX for cryo-EM and crystallography. Acta Crystallogr. D Struct. Biol. 74, 531–544 (2018).

52. Chou, F.-C., Sripakdeevong, P., Dibrov, S. M., Hermann, T. & Das, R. Correcting pervasive errors in RNA crystallography through enumerative structure prediction. Nat. Methods 10, 74–76 (2013).

53. Williams, C. J. et al. MolProbity: More and better reference data for improved all-atom structure validation. Protein Sci. 27, 293–315 (2018).

54. Afonine, P. V. et al. New tools for the analysis and validation of cryo-EM maps and atomic models. Acta Crystallogr. D Struct. Biol. 74, 814–840 (2018).

55. Richardson, J. S. et al. RNA backbone: consensus all-angle conformers and modular string nomenclature (an RNA Ontology Consortium contribution). RNA 14, 465–481 (2008).

56. Meng, E. C. et al. UCSF ChimeraX: Tools for structure building and analysis. Protein Sci. 32, e4792 (2023).

57. Nawrocki, E. P. & Eddy, S. R. Infernal 1.1: 100-fold faster RNA homology searches. Bioinformatics 29, 2933–2935 (2013).

58. Edgar, R. C. MUSCLE: a multiple sequence alignment method with reduced time and space complexity. BMC Bioinformatics 5, 113 (2004).

59. Ogurtsov, A. Y., Roytberg, M. A., Shabalina, S. A. & Kondrashov, A. S. OWEN: aligning long collinear regions of genomes. Bioinformatics 18, 1703–1704 (2002).

60. Kumar, S., Stecher, G., Li, M., Knyaz, C. & Tamura, K. MEGA X: Molecular Evolutionary Genetics Analysis across computing platforms. Mol. Biol. Evol. 35, 1547–1549 (2018).

61. Spiridonov, A. N. & Shabalina, S. A. Deciphering structural selective constraints: A comparative evolutionary analysis of RNA hairpin structures. in *Lecture Notes in Computer Science* 196–208 (Springer Nature Switzerland, Cham, 2024).

62. Bernhart, S. H., Hofacker, I. L., Will, S., Gruber, A. R. & Stadler, P. F. RNAalifold: improved consensus structure prediction for RNA alignments. BMC Bioinformatics 9, 474 (2008).

63. Ogurtsov, A. Y., Shabalina, S. A., Kondrashov, A. S. & Roytberg, M. A. Analysis of internal loops within the RNA secondary structure in almost quadratic time. Bioinformatics 22, 1317–1324 (2006).

64. Kondrashov, A. S. & Shabalina, S. A. Classification of common conserved sequences in mammalian intergenic regions. Hum. Mol. Genet. 11, 669–674 (2002).

65. Rivas, E., Clements, J. & Eddy, S. R. A statistical test for conserved RNA structure shows lack of evidence for structure in lncRNAs. Nat. Methods 14, 45–48 (2017).

## References

1. Rosenthal, P. B. & Henderson, R. Optimal Determination of Particle Orientation, Absolute Hand, and Contrast Loss in Single-particle Electron Cryomicroscopy. Journal of Molecular Biology 333, 721–745 (2003).

2. Pintilie, G. et al. Measurement of atom resolvability in cryo-EM maps with Q-scores. Nat. Methods 17, 328–334 (2020).

3. Hirano, S. et al. Structure of the OMEGA nickase IsrB in complex with ωRNA and target DNA. Nature 610, 575–581 (2022).

4. Young, G. et al. Quantitative mass imaging of single biological macromolecules. Science 360, 423–427 (2018).

5. Soares, L. W., King, C. G., Fernando, C. M., Roth, A. & Breaker, R. R. Genetic disruption of the bacterial *raiA* motif noncoding RNA causes defects in sporulation and aggregation. Proc. Natl. Acad. Sci. U. S. A. 121, e2318008121 (2024).

6. Nölling, J. et al. Genome sequence and comparative analysis of the solvent-producing bacterium Clostridium acetobutylicum. J. Bacteriol. 183, 4823–4838 (2001).

8. Weinberg, Z. et al. Detection of 224 candidate structured RNAs by comparative analysis of specific subsets of intergenic regions. Nucleic Acids Res. 45, 10811–10823 (2017).

9. Hess, M. et al. Metagenomic discovery of biomass-degrading genes and genomes from cow rumen. Science 331, 463–467 (2011).

10. Weinberg, Z., Perreault, J., Meyer, M. M. & Breaker, R. R. Exceptional structured noncoding RNAs revealed by bacterial metagenome analysis. Nature 462, 656–659 (2009).

11. 11. Yooseph, S., et al. The Sorcerer II Global Ocean Sampling expedition: expanding the universe of protein families. PLoS Biol. 5, e16 (2007).

